# WaterFlow: Prediction of Ordered Water Molecule Positions on Protein Structures

**DOI:** 10.64898/2026.08.26.747373

**Authors:** Vratin Srivastava, Huanghao Mai, Marcus Collins, James Holton, Michael Wall, Stephanie A. Wankowicz

## Abstract

Ordered water molecules mediate many protein functions including stability, ligand binding, and catalysis. Predicting their positions with sub-angstrom accuracy would support protein design, binding affinity prediction, and automated model building in X-ray crystallography and cryo-EM. However, water molecule prediction lags behind protein and other molecule structure predictions. We introduce WaterFlow, a flow-matching-based generator model and confidence model for predicting the positions of ordered water molecules in protein structures. WaterFlow outperforms the existing state of the art at every precision level. We demonstrate that this model not only predicts ground truth modeled water molecules, including those around ligands, but also fits the underlying experimental data well, and therefore proposes that it may be used for both prediction and modeling water molecules. We demonstrate that WaterFlow’s novel predictions are often associated with positive electron difference density, meaning the model places water molecules at sites the original structure depositions omitted. We use this improved model to address the data constraint. By mapping the Pareto front of achievable accuracy of water molecule prediction, alongside analysis of different training data schemas, we quantified the tradeoff between data quantity and data quality, demonstrating the diversity of high quality structures is limiting the results possible. Overall, WaterFlow predicts ordered water to serve as a solvent module for structure-based drug design, and predicted structures, as well as for water molecule placement during crystallographic refinement.

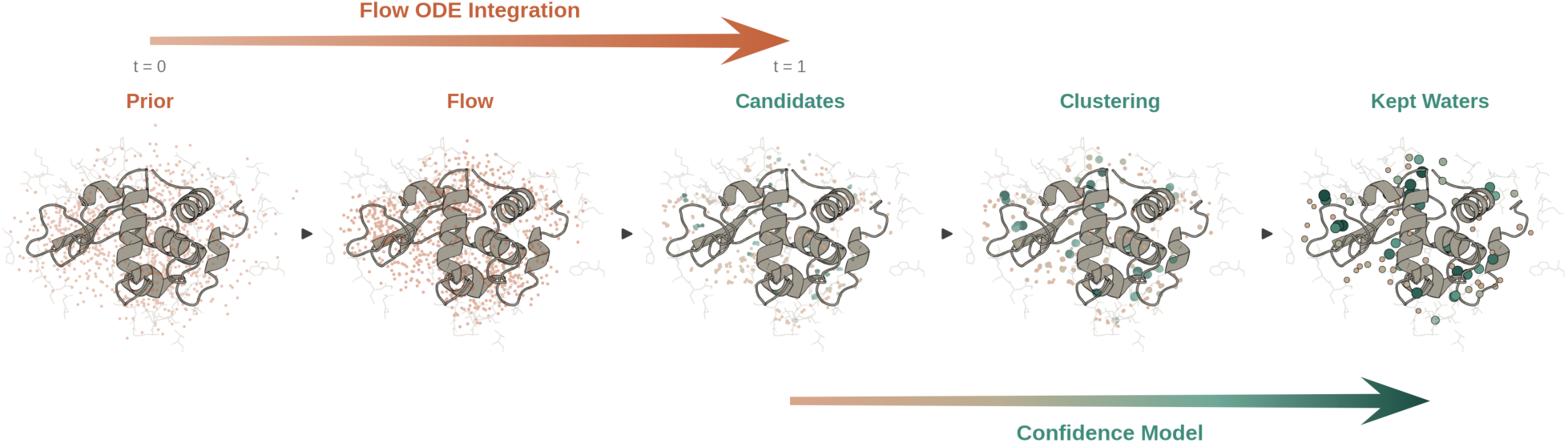

## Introduction

Water is essential for life. Most biological processes occur in aqueous environments, and the interactions between macromolecules and water are critical to every aspect of biology, including protein folding, stability, binding affinity, and catalytic efficiency [1–7]. Water molecules often participate in bridging interactions decisive for stability and binding affinity [8,9]. Water also directly participates in catalysis, both supporting the geometry of the active site and acting as a reactant, as in hydrolysis [10,11]. Interpreting the structural model of a macromolecule in light of its function or design often depends on accurate water placement. Sub-angstrom accuracy in predicted water molecule positions is therefore likely required to predict or design properties such as binding free energies [12–16], catalytic functions [17], and protein stability [18–20].

However, accurately predicting the water molecules positions remains challenging. This limits our ability to use this information for biological insights or design. This is in contrast to protein structure prediction which now achieves near-experimental accuracy for static structures [21–23]. The prediction of protein-small molecules and other macromolecules still lag behind proteins, but are rapidly becoming more precise [24–26]. Water molecule prediction has kept pace with neither. Most work on placing water molecules relies on physics-based models grounded in molecular dynamics and solvation thermodynamics [27,28]. These approaches are limited in speed, and their accuracy depends on the force field and solvation model used [29,30].

More recently, the prediction of water molecule placement has been approached with deep learning. HydraProt and GalaxyWater-CNN used a 3D convolutional neural network and showed that a learned model could achieve better water coverage than traditional physics-based approaches [29,31]. The current state of the art is SuperWater, a generative model that couples a score-based diffusion process with equivariant graph neural networks [32]. Against HydraProt, SuperWater recovers more water at every precision level. However, these predictions are still limited at high precision, where SuperWater reaches only roughly 27% recall. This ceiling reflects two possible constraints, the capacity of the model or by the quality of the data the model is trained on.

A predictive model of water molecule positions requires training on experimentally determined coordinates. These come primarily from X-ray crystallographic structures, which resolve protein and water molecule positions at the highest resolution [31,33]. Water constitutes approximately 50% of the volume of a typical protein crystal [34]. But, the majority of water molecules are disordered and are implicitly accounted for as bulk solvent. Water molecules can only be explicitly modeled when they are sufficiently ordered, typically due to hydrogen bonding with macromolecules, small molecules, or other water molecules [35–37]. These ordered water molecules are what are used for training prediction algorithms, but this data is very noisy as crystallographic models show substantial variance in which water molecules are modeled explicitly [38,39].

This variability has two sources. The first is physical difference between experiments, including temperature, unit cell size, pH, and pressure [40–44]. Second, the modeling process itself. This includes peak-picking algorithms during refinement [45,46] or manual selection using visualization software such as Coot [47]. Some of this modeling is done in real space. Reconstructing real-space density requires phase information that is not measured experimentally, so phases are estimated from the previous cycle’s model and the resulting density is biased toward that model. Map accuracy therefore depends on what has already been built, and any errors propagate forward [48]. Unmodeled density can arise from phase error, but it can also represent real signals from unmodeled water molecules, ligands, or from alternative locations of protein atoms [39,49]. These compounding sources of variation are borne out empirically. Water molecules in deposited biomolecular structures often fit the experimental data poorly and can be idiosyncratic to a single protein structure [50,51]. This variation increases at lower resolution, with the number of water molecules placed dropping sharply below 2.0 Å resolution and rarely modeled beyond 2.5 Å [35]. Reproducible water molecules that are physically present but absent from the deposited model are scored as a false negative, which bounds recall. Water that is present in one structure but missing from the specific structure used for evaluation is scored as a false positive, which bounds precision. Together these problems present a substantial obstacle for training a predictive model on deposited water molecule positions.

Here, we introduce WaterFlow, a flow-based model generator and confidence model for predicting the positions of ordered water molecules in protein structures. WaterFlow outperforms SuperWater at every precision level, establishing a new state of the art model for predicting water molecules. We demonstrate that this model not only predicts ground truth modeled water molecules, including those around ligands, but also fits the underlying experimental data well, proposing it may be used for both prediction and modeling water molecules. We use this improved model to address the data constraint. By mapping the Pareto front of achievable accuracy of water molecule prediction, alongside analysis of different training data schemas, we quantified the tradeoff between data quantity and data quality, demonstrating the diversity of high quality training data is limiting the results possible.

## Results

### Model Overview

#### Encoder, Generator, and Confidence Model

We developed WaterFlow (https://github.com/diff-use/WaterFlow) to predict the positions of ordered water molecules around proteins, taking into consideration the crystallographic environment most protein structures are derived from. WaterFlow consists of a candidate generator and a confidence model which scores the generated candidates. The two stages have the same core geometric vector perceptron (GVP) [52] architecture (**Fig. 1A**) which updates the node embeddings of a protein-water graph. We use ESM3 [53] residue embeddings as input features, broadcast to individual atom nodes. The generator is trained using a flow matching objective [54–56] to predict a linear vector field which connects a prior distribution with expected water positions from the graph embeddings. At inference time noisy water positions are sampled from a prior and the model predicts the vector displacements to the final positions. The confidence model encoder is initialized with the generator encoder weights, and encodes a graph with the protein and predicted water molecules from the generator. The confidence model is a regressor trained on a target which is ∼1 if the predicted water is within 0.5 Å of a ground truth water, and decays to zero at ∼1.5 Å. Further details are in the **Supplementary Methods.**

**Figure 1.**
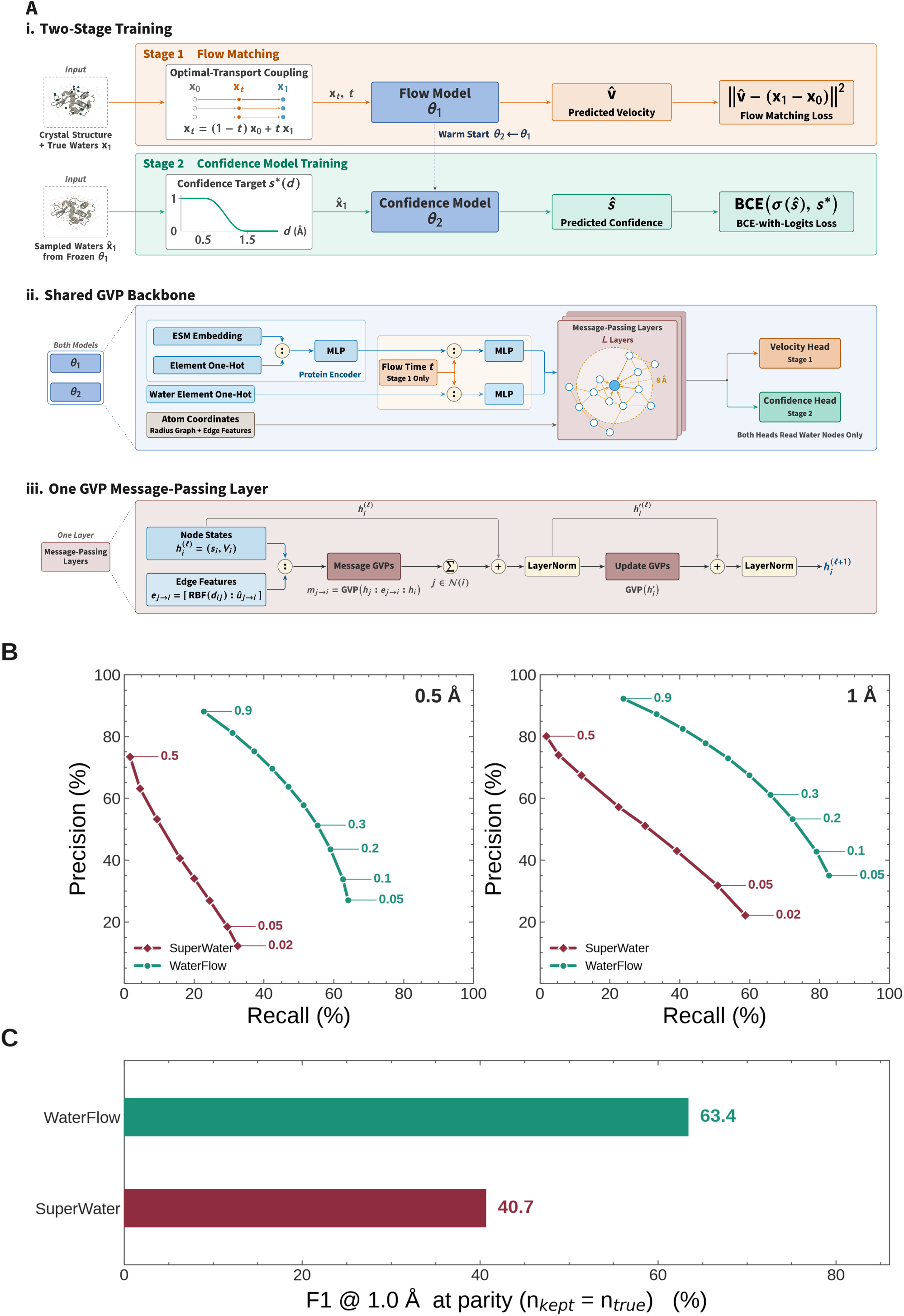
Model architecture and precision/recall of WaterFlow compared to SuperWater. **A.** WaterFlow architecture and training. (i) Training proceeds in two stages. In the first, a flow-matching generator learns a velocity field that transports protein-anchored prior samples onto crystallographic water positions, with a per-structure optimal-transport assignment linking interpolated and target positions. In the second, the generator is frozen and its sampled candidates train a confidence model against a distance-based target derived from crystallographic water molecules. (ii) Protein features, water coordinates, and a randomly sampled flow-matching time point are encoded and passed through L GVP message-passing layers acting on protein–protein and protein–water edges, which output per-water velocities or confidence scores. (iii) Each GVP layer builds source-to-destination messages from node states and distance and direction edge features, aggregates them over neighbors, and applies residual, layer-normalized message and node updates. **B**. Precision and recall for WaterFlow and re-trained SuperWater at 0.5 Å and 1.0 Å cutoffs, swept across confidence thresholds on our holdout dataset (n = 698). Confidence thresholds are swept over each model’s supported range. This is [0.05, 0.5] for SuperWater and [0.05, 0.9] for WaterFlow; see further details in **Methods**. The curves should be compared by precision and recall rather than by threshold value. **C**. Evaluation of the F1 score at 1.0 Å for WaterFlow and re-trained SuperWater models. The F1 is computed at parity, defined by the number of water molecules in the deposited structure, to enable a realistic comparison of performance in ideal settings.

#### Dataset

WaterFlow is trained on structures from PDB-REDO [45,57]. We built two datasets that trade structural diversity against the quality of water molecule placements. We first selected structures solved by X-ray crystallography containing 50-500 residues in polypeptide chains. The **LARGE** dataset includes structures with ≤ 3.0 Å resolution and R-free of ≤0.20, yielding 67,272 structures. **SMALL** includes structures with resolution ≤ 2.0 Å and R-free of ≤0.25, 22,318 total. As we filtered the R-free values from the RCSB but used PDB-REDO structures, some structures in our dataset had R-free above our cut off. Regardless, the **SMALL** dataset had better R-free (median 0.172, range 0.089–0.572) compared to the **LARGE** dataset (median 0.201, range 0.089-0.602). Full statistics for both datasets are in **Table 1** and **Supplementary Figures 1 and 7**.

**Table 1:** Dataset curation statistics for LARGE and SMALL.

|  | LARGE | SMALL |
| --- | --- | --- |
| Resolution | $\leq 3.0$ Å | $\leq 2.0$ Å |
| Number of Residues Per Structure | 50–500 | 50–500 |
| Number of Structures | 67,272 | 22,318 |
| Number of Sequence-Structure Clusters | 15,573 | 5,424 |
| Median Resolution | 1.73 Å | 1.44 Å |
| Median R-free | 0.201 | 0.172 |
| Median Number of Deposited Water Molecules per Structure | 182 | 250 |
| Structures with valid graph after filtering | 63,262 (94.0%) | 22,255 (99.7%) |
| Median Water Molecules Retained after EDIA/B-factor Filtering | 162 | 222 |

To prevent train–test leakage we cluster each dataset first by sequence using MMseqs2 [58] using a cutoff of 50% sequence coverage, then by structure using Foldseek [59], using a cutoff of min(qTM, tTM) ≥ 0.50 (where q- and t- indicate query or target length normalization, respectively). This procedure yielded 15,573 clusters in the LARGE dataset and 5,424 clusters in the SMALL dataset. Whole clusters are then assigned to training, validation, and test sets by structure count. SMALL dataset clusters are nested within LARGE’s clusters, so its hold-out sets are leak-free for either training dataset.

Additional filters based on B-factors and EDIA (Electron Density score for Individual Atoms, [60]) were also applied. In crystallography, the B-factor of an atom approximates the thermal motion of the atom. It correlates with the height of the electron density peak at the oxygen atom in water [61] and is often treated as a confidence metric for the atomic model [62–64]. The EDIA [60] evaluates how well an atom’s placement is supported by the map using a weighted sum of electron density near the atom (**Supplementary Figure 2**). An EDIA above 0.8 indicates a strong support by the map whereas below 0.4 suggests inadequate or misfit to the density map [60]. Further details are described in the **Supplementary Methods.**

#### Evaluation Dataset

All models are evaluated on a single holdout set disjoint from the training set, and the validation set used to select best model checkpoints.The SMALL dataset is, by construction, nested within the LARGE dataset; the same holdout structures therefore are unseen by every model, regardless of the training dataset. We obtain 698 structures for our evaluation dataset, which are used for all evaluations in the subsequent sections unless stated otherwise. These structures have a median resolution of 1.90 Å, median R-free of 0.212, and a median water to residue ratio of 0.51. Structure-level data on both the evaluation and training data is available in the supplementary data.

#### Inference

At inference two networks are applied in sequence (**Fig. 1A**). The generator samples several-fold more candidate water molecules than are expected for the structure, and we score each with the confidence model. Candidates below an acceptance confidence or density threshold, as defined by the user, are discarded. We follow SuperWater [33] to cluster the accepted candidates greedily within an oxygen van der Waals radius (1.52 Å). From each cluster we compute a confidence-weighted mean center; this can result in cluster centers within a water molecule radius of each other, in which case we take the highest scoring cluster and discard the others. The output is a discrete, confidence-ranked set of predicted water molecules. The thresholds control how many water molecules are returned (**Fig. 1B**).

#### Evaluation Metrics

In this context we define precision@d as the fraction of predicted water molecules lying within distance *d* of some deposited water. Similarly recall@d is the fraction of target water molecules within *d* of some predicted water. We report these metrics at both acceptance radii *d* = 0.5 and 1.0 Å, averaged over the holdout dataset. Note that the matching is not necessarily one-to-one. In general precision and recall are computed at some confidence threshold, or alternatively at some fixed ratio of predicted to target water molecules. We summarize precision and recall by the geometric mean of precision and recall F1 = 2PR/(P+R), at parity, i.e., the point where the numbers of target (deposited modeled water molecules) and predicted water molecules are the same.

#### WaterFlow More Accurately Predicts Water Molecules Compared with Existing State of the Art

To assess the improvements of WaterFlow over the existing state of the art, SuperWater [32], we re-trained SuperWater using the LARGE and SMALL dataset evaluating on the same holdout as WaterFlow. Precision-recall curves generated by sweeping the minimum confidence threshold are plotted for each model in **Fig 1B**.

WaterFlow has much better recall at any given precision, and vice versa, compared to SuperWater. At 1.0 Å acceptance radius, WaterFlow achieves a maximum precision and recall of 0.92 and 0.83 against SuperWater’s 0.8 and 0.59. Both models lose precision and recall at the stricter 0.5 Å acceptance radius, however, WaterFlow’s maximum precision/recall only fell to 0.88/0.64, while SuperWater’s precision/recall fell to 0.73/0.33. For both models, we report numbers obtained from the best performing checkpoints. For WaterFlow this was on the LARGE dataset, and for SuperWater the best results were obtained on the SMALL dataset.

We then examined the trade-off at parity, which we defined as the operating point at which the number of predicted water molecules equals the number of target water molecules (**Fig. 1C**). At the parity operating point and a 1.0 Å acceptance radius, WaterFlow reaches precision 0.64 and recall 0.63 (F1 = 0.63), against 0.42 and 0.41 for SuperWater (F1 = 0.41). Tightening the radius to 0.5 Å widens the parity F1 gap (0.54 for WaterFlow vs 0.26 for SuperWater), demonstrating that WaterFlow’s largest advantage is at sub-angstrom localization. We also report numbers obtained from the raw generators from both WaterFlow and SuperWater in **Supplementary Table 1**, where we examine the performance of the generators as one-shot predictors of water placement without confidence/ranking heads.

We note that the precision and recall we report for the retrained SuperWater curves are lower than originally reported [33]. We attribute this to a difference in dataset splitting. SuperWater was trained and evaluated on random train/validation/test splits (8:1:1). The authors’ stated rationale was that structurally similar proteins can adopt substantially different water arrangements, as measured by the root mean squared distance (RMSD) between the coordinates of water molecules for a given set of homologs. However, as we will demonstrate in the next section, homologous proteins often have a high number of overlapping water sites. To decrease the potential effects of memorization, we modified the splitting method when re-training SuperWater, leading to a decrease in the precision and recall.

#### Crystallographic Water Placement Pareto Front

While WaterFlow substantially improves over existing water prediction methods, its predictions are still below what may be needed for demanding biophysical or biochemical applications. We therefore explored the limitations of the training data itself. We and others have trained their model conditioned on proteins and ligands [29,31,32]. However, temperature, pH, pressure, and crystallization conditions all affect water molecule placement [40–43]. If deposited water molecules differ substantially between similar structures under different conditions, that variation is irreducible from the model’s perspective and will limit the estimated accuracy. To quantify these bounds, we conducted a retrospective analysis to quantify the variance in deposited water molecules among similar structures.

We first examined deposited, modeled water molecules among proteins that are similar by sequence and structure and are isomorphous, meaning they have the same unit cell and space group. We began with a small curated dataset (n = 65) of hen egg white lysozyme (HEWL) structures with R-free better than 25% and resolution better than 2.0 Å (**Supplementary Table 2**). All structures had identical amino acid sequences and no ligand at the binding site. We selected the highest resolution structure as the reference (PDB: 7DER). All other structures had less than 0.42 Å pairwise C_ɑ_ RMSD to the reference structure after alignment. We then clustered water molecules from all structures with HDBSCAN [65,66], with hyperparameters grid-searched and ranked by DBCV [67] and cluster stability (**Fig. 2A; Methods**). We defined the consensus water molecules as those which belong to clusters having occupancy above 0.3, meaning at least 30% of structures in the dataset have a water molecule explicitly modeled within 1.0 Å.

**Figure 2.**
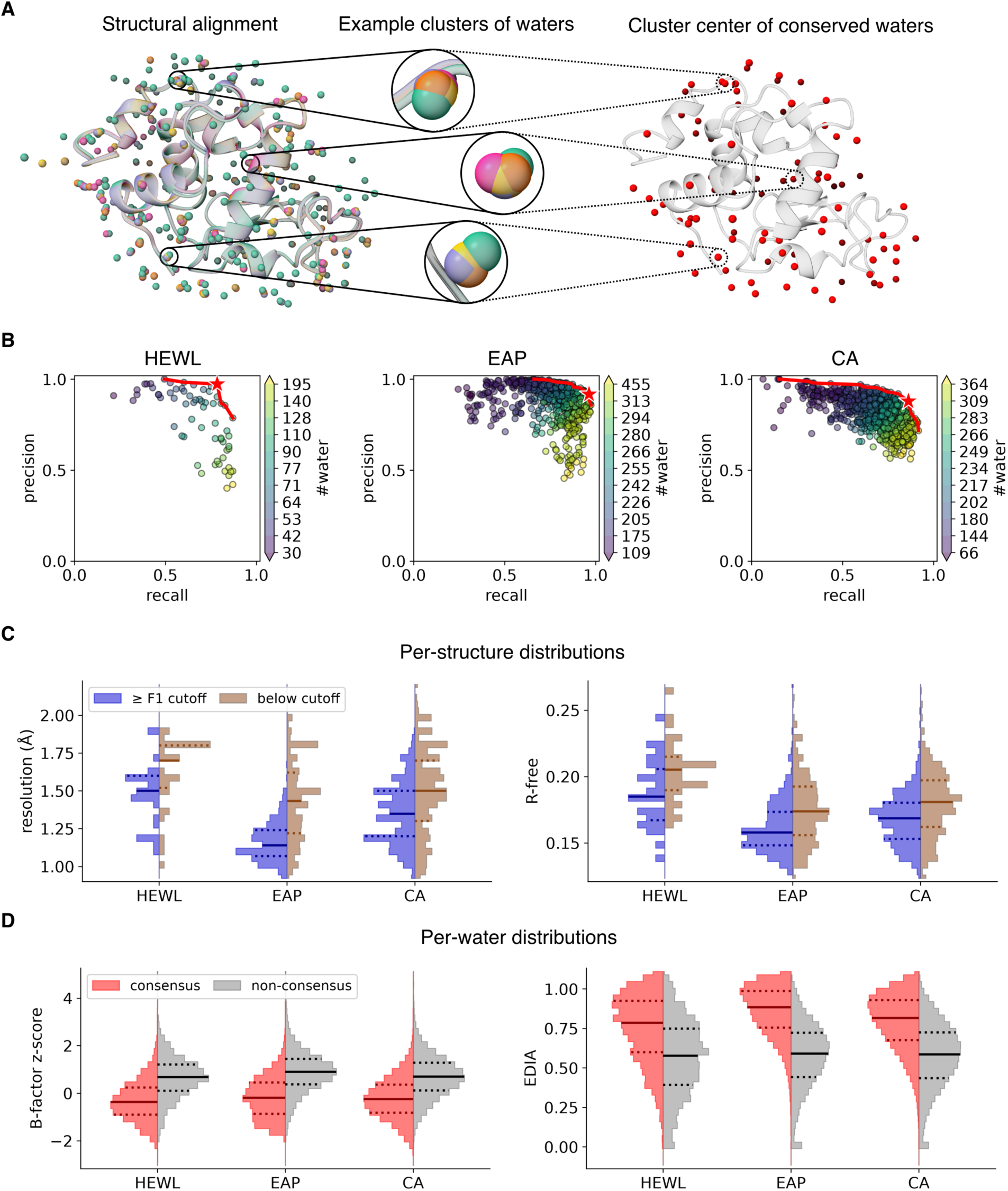
Water molecule placements across isomorphous structures. **A.** Schematic of water molecule clustering analysis. For a given dataset, we identify the conserved water sites by aligning the protein structures and then performing clustering analysis on the deposited water molecules. The center of a highly occupied cluster of water molecules marks a consensus water position in this dataset. **B.** The precision and recall scoring of each PDB entry’s water against the consensus sites using a distance cutoff = 1.0 Å. Each PDB is colored by the number of water molecules it contains. The solid red line traces the empirical Pareto front, with the knee identified by the maximum F1 and marked with a red star. **C.** Distribution of the resolution and the R-free (from PDB-REDO) of the PDBs in each dataset. Within each dataset, the PDBs are split into two halves by their F1 score of the deposited water molecules against the consensus water molecules. Each distribution’s median value and IQR are marked by solid and dotted lines respectively. **D.** Distribution of the B-factor z-score and the EDIA (queried from PDB-REDO) of the deposited water molecules in each dataset. Within each dataset, the water molecules are split by whether they are consensus water molecules or not. **C** and **D** clip the visualization at 99.9 percentile of the values. Results of **B-D** using a distance cutoff = 0.5 Å are shown in **Supplementary Figure 3**.

The best clustering result defined 69.9% of water molecules as consensus, with values ranging from 63.2% to 70.4% across hyperparameter settings. This means that the remaining 30.1% of water molecules appear in fewer than 30% of structures. We also performed the same clustering on two larger datasets of different protein families, carbonic anhydrases (CA, n = 937; **Supplementary Table 3**) and endothiapepsin (EAP, n = 907; **Supplementary Table 4**). Both datasets contain isomorphous structures with R-free better than 25% but vary in terms of bound ligands and mutants. In the EAP dataset, the fraction of conserved water molecules ranges from 83.6% to 86.5%, with the best clustering result assigning 86.5% water molecules as conserved. The CA dataset varies more on the sequence identities and contains more lower-resolution structures (up to 3.0 Å). The fraction of conserved water ranges from 77.8% to 79.1%, with the best clustering result assigning 78.5% water molecules as conserved. EAP and CA likely differ in the percentage of water molecules found in consensus positions because the EAP dataset is more homogeneous and made of higher resolution.

Treating the centers of the conserved clusters as the “ground truth” water molecule positions, we computed precision, recall, and F1 score of water molecules for each deposited structure. All three datasets exhibit a clear trade-off between precision and recall as a function of the number of water molecules modeled (**Fig. 2B; Supplementary Table 5**). For the HEWL dataset, structures with the fewest number of water molecules modeled have an average precision of 0.94 and a recall of 0.36, whereas structures with the highest number of water molecules modeled have a precision of 0.47 and a recall of 0.85. As expected, with more water molecules modeled, recall improves at the cost of lower precision, and vice versa. For the two larger datasets, precision/recall ranges from 0.93/0.47 to 0.70/0.89 for EAP and 0.92/0.36 to 0.68/0.84 for CA. We traced the Pareto front for precision and recall across all datasets, with the best point (Pareto knee) identified by the maximum F1 (HEWL: precision/recall/F1: 0.97/0.78/0.87; EAP: 0.92/0.96/0.94; CA: 0.88/0.86/0.87). These numbers all fall slightly with a cutoff distance for 0.5 Å instead of 1.0 Å, but follow a similar pattern (**Supplementary Figure 3**; **Supplementary Table 5**). Overall, the theoretical F1 scores demonstrate there is still significant room for improvement with WaterFlow.

Next, we examined if quality metrics both on a per-structure and per-water level track water consensus. On a per-structure level, we analyzed how resolution and R-free related to the consensus “ground truth water molecules”. We split each dataset (HEWL, EAP, CA) at its median F1 score and performed Mann–Whitney U tests to see if the two halves differ significantly. In all three datasets, the half with higher F1 scores has better resolution and lower R-free (**Fig. 2C**; **Table 2**). Better resolution supports a better structural model where more ordered water molecules are modeled, which raises the recall and drives the F1 difference (**Supplementary Figure 4**).

**Table 2:** Distribution of the resolution and R-free of structures above versus below the median F1 score (HEWL: 0.71, EAP: 0.85, CA: 0.74) against the consensus set of water molecules for each dataset. The median value and IQR are reported along with the rank- biserial correlation (r) and Mann–Whitney test p-value (p).

| metric | dataset | above |  | below |  | n_above | n_below | r | p |
| --- | --- | --- | --- | --- | --- | --- | --- | --- | --- |
|  |  | median | iqr | median | iqr |  |  |  |  |
| resolution (Å) | HEWL | 1.5 | (1.37, 1.6) | 1.7 | (1.52, 1.8) | 31 | 34 | -0.41 | $4.88 \times 10^{-3}$ |
| | EAP | 1.14 | (1.07, 1.24) | 1.43 | (1.22, 1.62) | 453 | 454 | -0.61 | $1.26 \times 10^{-57}$ |
| | CA | 1.35 | (1.2, 1.5) | 1.5 | (1.3, 1.7) | 468 | 469 | -0.32 | $8.82 \times 10^{-18}$ |
| R-free% | HEWL | 18.48 | (16.71, 20.55) | 20.51 | (18.97, 21.49) | 31 | 34 | -0.39 | $7.82 \times 10^{-3}$ |
| | EAP | 15.78 | (14.81, 17.33) | 17.37 | (15.58, 19.25) | 453 | 454 | -0.32 | $3.77 \times 10^{-17}$ |
| | CA | 16.86 | (15.3, 18.03) | 18.09 | (16.21, 19.71) | 468 | 469 | -0.3 | $7.1 \times 10^{-16}$ |

We also compared the distribution of normalized B-factors and EDIA between consensus and non-consensus water molecules, with the hypothesis that consensus water molecules would be better supported by experimental data. Because water molecules from the same structure are not independent samples, we performed 500 structure-level bootstrapping to compute the 95% confidence interval (CI) of the estimated rank biserial correlation. For comparison across structures we compute the z-scores for B-factors of water molecules within each structure. The distribution of B-factor z-score for consensus water has a similar shape and spread to that of non-consensus water, but the median is shifted towards more negative values, indicating lower B-factors (**Fig. 2D**; **Table 3**). In all three datasets, the consensus water molecules’s EDIA is skewed towards higher values (**Fig. 2D; Table 3**). Removing water molecules with EDIA below 0.4 would eliminate 20-26% of non-consensus water molecules but only 1-8% of consensus water molecules. Strong real-space density is often correlated with low thermal motion, and enables less ambiguity when placing water molecules during structural refinement.

**Table 3:** Distribution of the B-factor z-score and the EDIA of consensus versus non-consensus water molecules in each dataset. The median value and IQR are reported along with the rank-biserial correlation (r) and its 95% confidence interval (CI) from bootstrapping.

| metric | dataset | consensus |  | nonconsensus |  | n_consensus | n_nonconsensus | r | CI |
| --- | --- | --- | --- | --- | --- | --- | --- | --- | --- |
|  |  | median | iqr | median | iqr |  |  |  |  |
| B-factor z-score | HEWL | -0.37 | (-0.9, 0.23) | 0.68 | (0.1, 1.21) | 4043 | 1745 | -0.57 | (-0.6, -0.53) |
|  | EAP | -0.19 | (-0.87, 0.45) | 0.91 | (0.37, 1.44) | 197290 | 30791 | -0.61 | (-0.62, -0.6) |
|  | CA | -0.25 | (-0.82, 0.36) | 0.7 | (0.11, 1.29) | 169187 | 46214 | -0.54 | (-0.55, -0.53) |
| EDIA | HEWL | 0.78 | (0.6, 0.92) | 0.58 | (0.39, 0.75) | 4043 | 1745 | 0.43 | (0.36, 0.49) |
|  | EAP | 0.88 | (0.76, 0.99) | 0.59 | (0.44, 0.72) | 197290 | 30791 | 0.72 | (0.71, 0.73) |
|  | CA | 0.82 | (0.67, 0.93) | 0.59 | (0.43, 0.72) | 169186 | 46214 | 0.57 | (0.56, 0.58) |

When using a distance cutoff = 0.5 Å (**Supplementary Figure 3**), the B-factor z-score and EDIA of water molecules still track their consensus status well (**Supplementary Table 6**). In comparison, the per-structure level metrics are more sensitive to the cutoff especially on the smallest dataset HEWL and become less effective (**Supplementary Table 7**). The direction of change on resolution and R-free between the half-dataset partitioned by median F1 nevertheless stayed the same. Overall, these two analyses support the idea that consensus water molecules have better underlying experimental data support and indicate that this may be a way to improve the training data for a water molecule prediction model.

#### Training data size affects prediction accuracy

Our analysis on the Pareto front of crystallographic water molecule placement showed that up to 30% of deposited water molecules are not reproduced across otherwise identical structures. We hypothesized that if that much of the training signal is unreliable, then restricting training to better-resolved and better-refined data should produce a more accurate model. We tested this by training WaterFlow on two datasets that trade quantity against quality. The SMALL dataset (n = 22,318) contains higher- resolution (≤ 2.0 Å), lower R-free (median: 0.172), compared to the LARGE dataset (n = 67,272; resolution: ≤ 3.0 Å; median R-free: 0.201). The structures in the SMALL dataset also contain more modeled water molecules (median of 1.01 v.0.77 water molecules per residue per structure; 0.90 v. 0.67 water molecules per residue after the per- water quality cutoffs, **Supplementary Figure 6**). However, the LARGE dataset has 3x the structures and 2.9x the cluster diversity.

To test the trade off of dataset size and quality, we retrained the full pipeline on periodically larger, cluster- stratified fractions of SMALL and LARGE, scoring every run on the same 698-structure holdout dataset (**Fig. 3A**). When the LARGE and SMALL dataset are of comparable sizes, the SMALL dataset achieves better F1 score, indicating that at comparable training dataset sizes, the higher quality data improve our predictions. However, as the training dataset gets larger, this gap narrows. Likely this is due to the fact that the small dataset contains around 3x fewer clusters (5,424 vs 15,573) so that above 10,000 examples the SMALL dataset is no longer adding diversity while the LARGE dataset still has additional clusters to sample. As we sample more deeply from the LARGE dataset, we see that its performance also saturates. This suggests that additional distinct high quality examples should improve the overall performance of our models.

**Figure 3.**
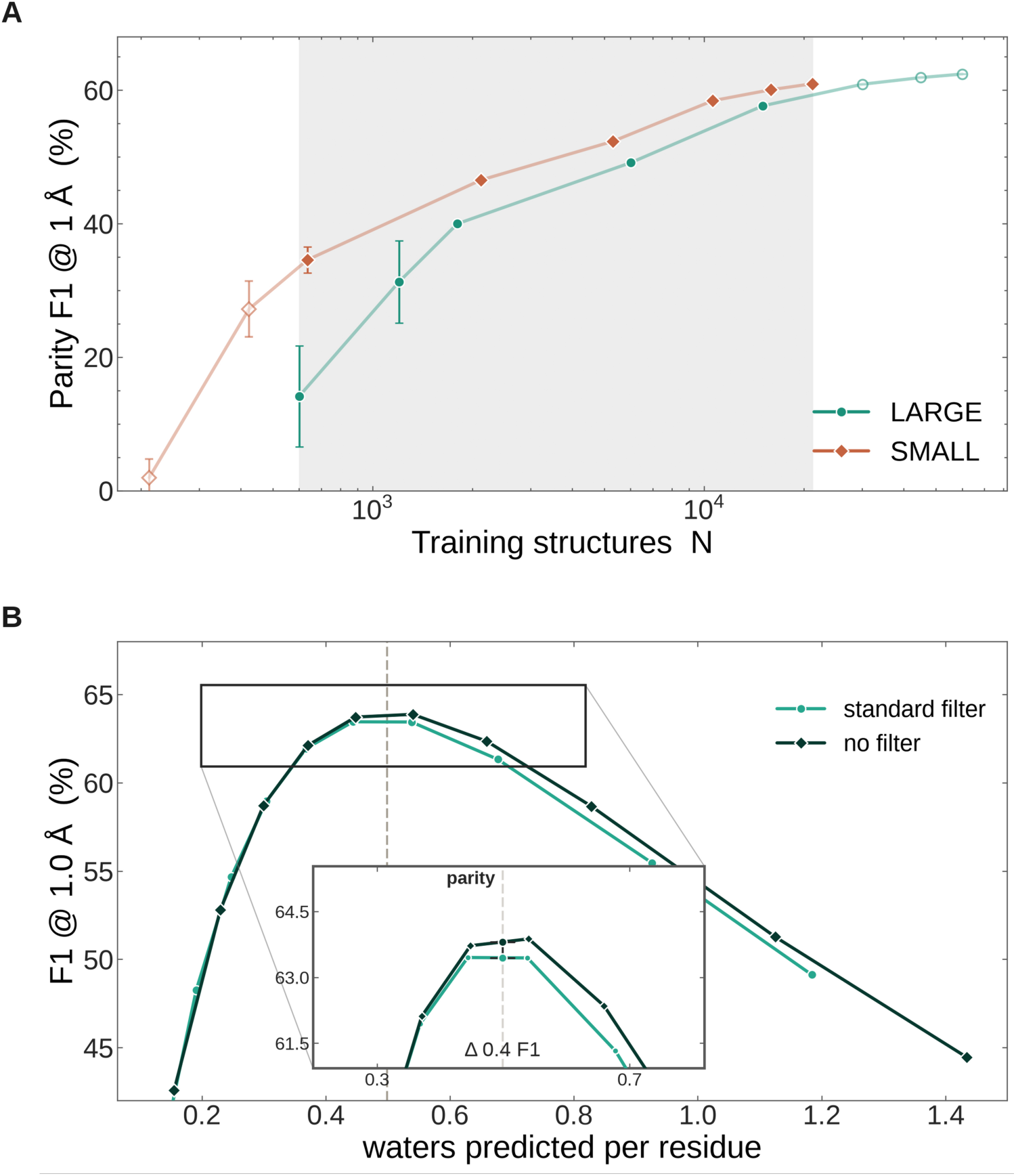
Training-set size scaling and data quality filtering. **A.** The F1 value at parity for 1.0 Å for water molecules in the holdout dataset (n=698) based on different training set sizes N for the LARGE (teal) and SMALL (brown) datasets. Training set fractions start at 1% and go to 100%. The shaded band (601–21,211 structures) indicates where N is the same for the two curves. The three smallest fractions of each dataset were replicated over three generator seeds and error bars are their standard deviation. The LARGE dataset passes SMALL’s best only by using more structures than the SMALL dataset has, reaching 63.4 at N = 60,116 against 61.0 for SMALL at 21,211. **B.** F1 at 1.0 Å for the LARGE-trained generators with the standard per-water filter (teal; EDIAm ≥ 0.4 and within-structure B-factor z ≤ 2.0) and with no per-water filter (dark green), on the 698 holdout dataset. The x-axis tracks the water molecules predicted per ASU residue regardless of threshold, and the F1 at that point, to separate the effect of the confidence threshold. The dashed line marks parity. Inset: the region either side of parity. The arms differ by 0.4 F1 at parity (63.4 filtered against 63.8 unfiltered) and by 0.50 F1 averaged across the matched window, always in favor of the unfiltered arm.

We then asked if the quality of the individual water molecules within a structure can help prediction accuracy. To test this, we removed water molecules with an EDIAm < 0.4 and normalized B-factor z-score > 2.0 (normalized over water molecules within the structure) from the LARGE dataset, dropping 14.1% of water molecules. We then re-trained WaterFlow with this filtered dataset and evaluated it on the same 698-structure holdout dataset. The model trained with unfiltered water molecules had a slightly better performance at a 1:1 ratio of predicted to target water molecules (F1 63.4 filtered v. F1 63.8 unfiltered; **Fig. 3B**). We speculate that these additional “lower quality” water molecules may have a slight regularizing effect on our models, but cannot be sure without more investigation. More likely, the diversity of structures in the LARGE dataset is what is improving accuracy there. Therefore, more diverse, high-quality protein structures are likely needed to improve water molecule predictions.

#### Encoding Symmetry Mates Improves Prediction Accuracy

Many ordered water molecules at the protein surface are coordinated by atoms across the crystal lattice. Indeed, water-mediated hydrogen bonds are characteristic of these crystal packing contacts [68,69] (**Fig. 4A**). To a model that sees only the asymmetric unit, the partners that coordinate such a water molecule are absent. In WaterFlow, we add these “symmetry mates” that pack against the asymmetric unit (ASU) to the input graph as additional protein nodes, giving the generator the complete local environment of each water rather. The symmetry expansion keeps whole residues, including ligands, ions, or cofactors, but not water molecules, with any atom within 8.0 Å of the ASU (**Methods**).

**Figure 4.**
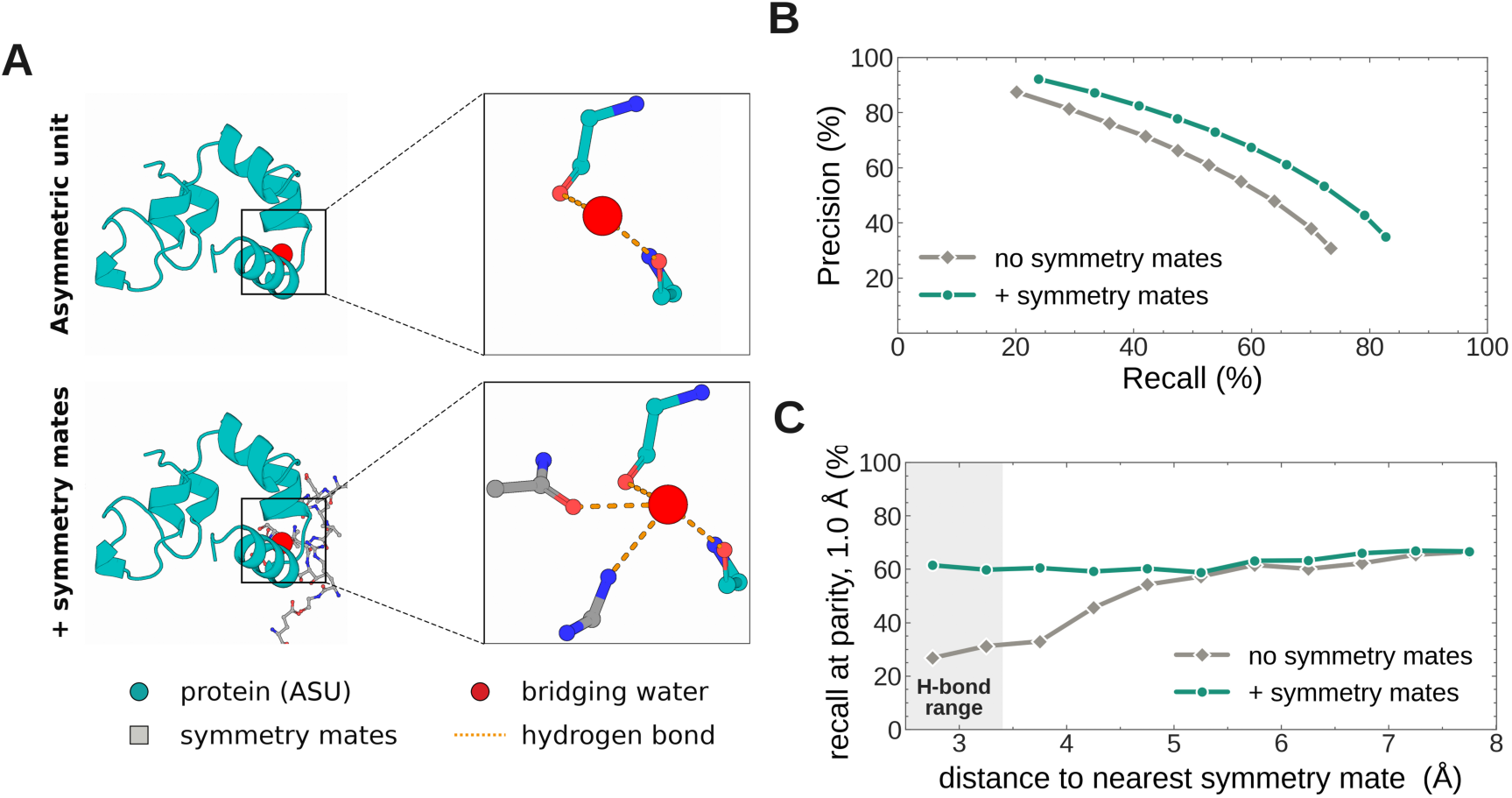
Adding symmetry mates as context during training and inference improves model performance. **A.** Water molecules at symmetry mates are coordinated by symmetry neighbors that are absent from the asymmetric unit. A representative bridging water (red) in PDB 3MYE, shown for the asymmetric unit alone (top) and with its symmetry mates added in grey (bottom). The blown up view shows water molecules within the H-bonding range (polar contacts ≤ 3.4 Å) with orange dashes. **B.** Precision/recall curve with or without symmetry mates. Both arms are trained on the LARGE dataset and differ only in the inclusion of crystallographic symmetry mates. Each point is a confidence threshold in the sweep (0.05-0.9). Symmetry mates raise precision by ∼0.1 and recall throughout the sweep. **C.** The symmetry mates model recovers more water molecules where a symmetry neighbor can coordinate it. Ground-truth water molecules (holdout, n = 698 structures) are binned by distance to the nearest symmetry mate atom. Each point groups all ground-truth water molecules whose nearest symmetry-mate atom falls in a distant bin, and the point on the curve represents the center of the bin. The y-axis is recall within 1.0 Å for the baseline (*default*, grey) and symmetry-mates- based (green) models, scored at parity. Each model’s threshold is set so it predicts as many water molecules as exist (equal budget), removing the confounder that the crystal-contacts model emits a denser cloud of candidates at the same sampling ratio due to higher number of residues. The shaded band indicates the range of hydrogen bonds to a symmetry mate (≤ 3.4 Å). Without symmetry mate information, only 28.8% of water molecules are recalled in this region, compared to 59.8% with symmetry mates. Beyond ∼5.0 Å, the impact of symmetry mates diminishes, however, the symmetry mate model always does slightly better on recall.

To isolate the contribution of including symmetry-related copies, we trained two models that differ only in whether symmetry mates are present in the input graph. When sampling water molecules from the flow model to score with the confidence model, with the symmetry mates, we sample more water molecules (∼1.5-2x) since we now have added context residues from the symmetry mates **(Supplementary Figure 8**). Beyond this, the additional context of the symmetry mates increases the confidence of many water molecules (**Supplementary Figure 9**). At parity (1:1 ratio of deposited to predicted water molecules) at 1.0 Å, including symmetry mates improves F1 by 0.06 (F1/precision/recall: with symmetry mates: 0.63/0.64/0.63 v. without symmetry mates: 0.57/0.57/0.57; **Fig. 4B**).

To test how this impacts different water molecule placements, we stratified every water molecule in the holdout dataset by its distance to the nearest symmetry mate atom. The gain in accuracy is sharply localized to the water molecules within the hydrogen-bonding range of symmetry mates (≤ 3.4 Å; ∼10% of all water molecules; **Fig. 4C**). For these water molecules, recall rose from 0.29 to 0.60, whereas for water molecules away from symmetry mates (> 3.4 Å), recall only moved from 0.62 to 0.67. Almost the entire gain at parity, is therefore attributable to the minority of water molecules whose hydrogen-bond network is likely completed by the lattice. Analyzing the confidence score assigned to a candidate water molecule within the hydrogen-bonding range of symmetry mate supports this hypothesis. The same water molecule generated with or without symmetry mates has a confidence score of 0.71 with symmetry mates compared to 0.004 without symmetry mates (**Supplementary Figure 9**).

#### WaterFlow Inference on Predicted Structures and Protein-Ligand Interactions

During inference, WaterFlow takes in a structure and returns a solvated structure. Which water molecules and the number of water molecules predicted on the returned solvated structures are determined by the confidence model. After the initial generation of candidates, a confidence model scores each candidate water molecule. By applying a confidence threshold, WaterFlow returns all water molecules with a confidence score above the specified threshold. The higher the threshold, the fewer number of water molecules are returned (**Fig 5B**).

**Figure 5.**
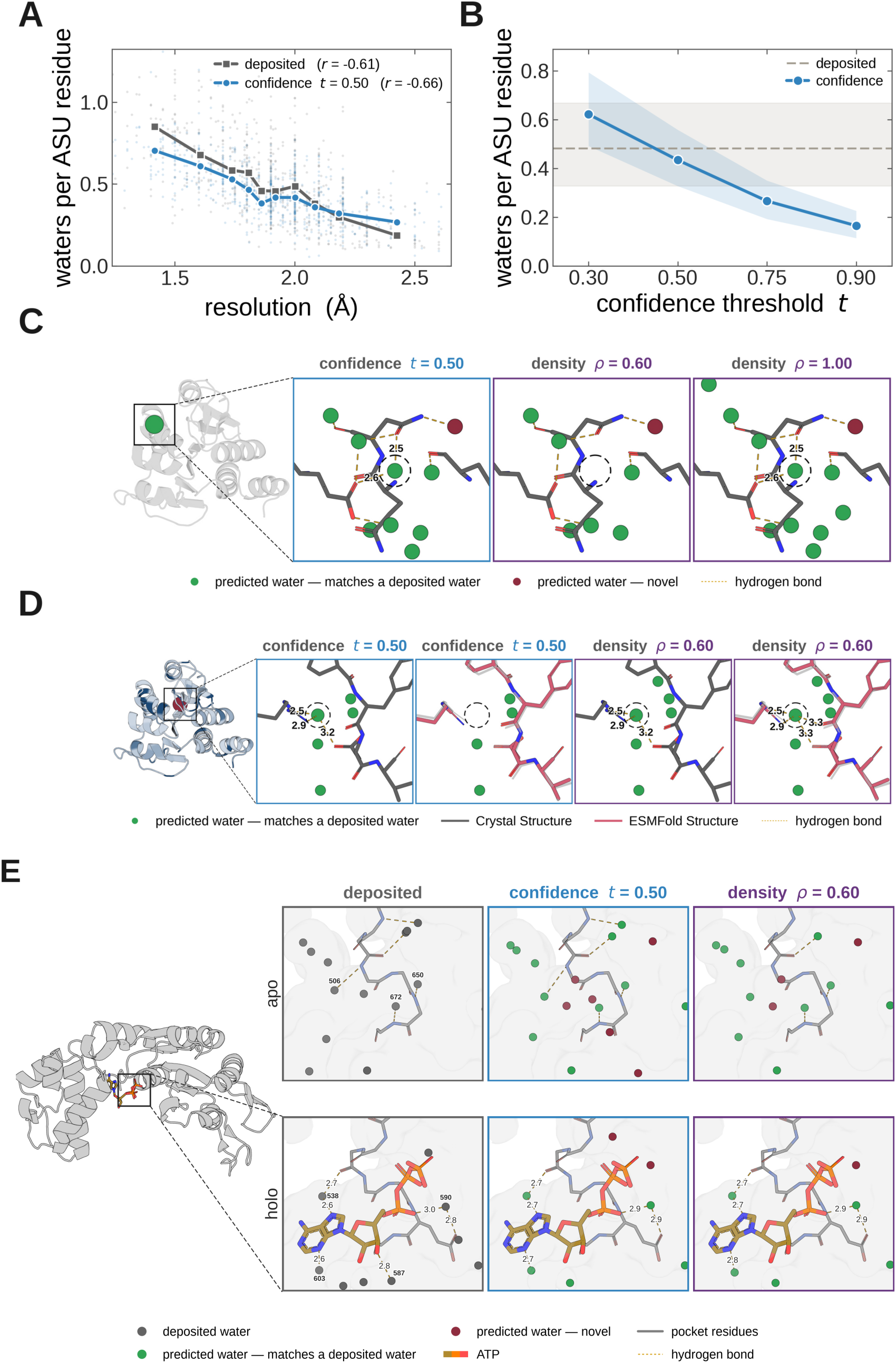
Using WaterFlow on predicted structures and with protein-ligand interactions. **A.** Water molecules per ASU residue versus resolution in the holdout dataset (n = 698). Grey: deposited water molecules. Blue: confidence model at t = 0.50. Faint points are individual structures, with joined markers are medians within deciles. Deposited water density (number of water molecules per structure) correlates with resolution (r = −0.61). Predicting water molecules with the confidence threshold also correlates with resolution (r = −0.66). **B.** Water molecules per ASU residue against the confidence threshold for the holdout dataset (n = 698), with different confidence thresholds. Lines are means, shaded bands the interquartile range across structures; the grey dashed line and band are the deposited median and IQR. **C.** Water molecules predicted on PDB 4RKW (1.50 Å). Green: predicted water within 1.0 Å of a deposited water. Crimson: novel water molecules. Orange dashes: potential bonds within H-bonding range (within 3.4 Å) to protein N or O. **D.** Left, the ESMFold mode overlaid with colored by per-atom displacement from the crystal, where dark blue is higher displacement. Green spheres show where the target water is recovered. Orange dashes are potential hydrogen bonds within 3.4 Å of the target water. **E.** Evaluating predicted water molecules at the ATP binding site of the apo/bound pair of PDBs 5EP1/5EP4 (1.5 Å). The inset boxes are the deposited water molecules (grey), the predicted water molecules at confidence threshold t=0.5 (blue) and density threshold ρ = 0.6 (violet). Grey spheres are the deposited water molecules, the green spheres are predicted water molecules within 1.0 Å of a deposited water, and the crimson spheres are un-matched, novel predicted water molecules. Dashed yellow lines are the potential hydrogen-bonds between water molecules and ligand/protein atoms. Bold lettering next to water molecules in the deposited panels are the water residue IDs.

Across our holdout dataset, when we applied a confidence threshold of t = 0.5, we observed that the number of predicted water molecules correlated with both the resolution and the average (Wilson) B-factor of the structure model (*R* = -0.66; **Fig. 5A; Supplementary Figure 5B**). These two correlations also exist in the deposited structures and modeled water molecules (*R* = -0.61; **Fig 5A; Supplementary Figure 5B**). Such a dependence is unlikely to be a property of the protein itself, but a reflection of the experimental data, with prior studies demonstrating the same trend [50]. To remove the correlation with resolution, we used the confidence score as a ranking, applying a ‘density threshold’ selecting the top scoring water molecules based on the number of residues in the structure. We repeated our precision/recall analysis, now sweeping density (ρ) instead of confidence (t), evaluated at a match radius of 1.0 Å. The curve for density thresholding sits below the confidence thresholding curve across all operating points, with a maximum precision of 0.879 at ρ = 0.1 and recall of 0.811 at ρ = 1.2. At a parity ratio of ρ = 0.5, density thresholding achieves precision of 0.595 and recall of 0.596 (F1 = 0.596) compared to a parity F1 of 0.634 for confidence thresholding (**Supplementary Figure 5A**). However, our density model accuracy still sits well above SuperWater [32].

To demonstrate how different inference settings (confidence v. density) place water molecules, we apply them to the 1.5 Å structure of DJ-1 (PDB 4RKW) [70] (**Fig 5C**). We tested a confidence threshold t = 0.5 (which gives us near optimal F1 balanced with high precision), and density thresholds ρ of 0.6 and 1.0. WaterFlow predicted 157 water molecules for t = 0.5, 112 for ρ = 0.6, and 186 for ρ = 1.0. **Supplementary Tables 8-11** show prediction statistics on 4RKW across the range of confidence and density thresholds, evaluated with both the symmetry-mates on and off models. Around the water molecule cluster formed by Leu77, Asn81, and Glu84, all three settings recover deposited waters within 1.0 Å and place an additional water within hydrogen bonding distance of the Asn81 side-chain amide (**Fig 5C**). One deposited water coordinated by Asn81 and Glu84 is absent at ρ = 0.6 but present at the other settings (center of each panel in **Fig 5C**), consistent with the larger number of waters predicted at higher ρ.

We next wanted to know how much a predicted structure might change the results relative to the crystal structure. We used ESMFold [71] to predict the structure of DJ-1 from the sequence of PDB 4RKW. Since we do not have any crystal information for the predicted structure, we use the model checkpoint trained without symmetry mates for inference. The predicted and deposited structures agree closely, with a Cα RMSD of 0.31 Å and an all-heavy-atom RMSD of 0.73 Å. We observed that the water molecule predictions on the ESMFold structure were worse than the crystal structure both in terms of precision and recall regardless of whether the confidence or density threshold are used (**Table 4**).

**Table 4:** The precision/recall on ground truth water molecules in PDB 4RKW generated from the crystallographic structure or EMSFold structure. For each baseline structure, we ran different inference modules (confidence or density) and included or removed symmetry mates.

| Input Structure | Method | Symmetry Mates | Candidate Water Molecules | Water Molecules Kept | Precision @ 1.0 Å | Recall @ 1.0 Å |
| --- | --- | --- | --- | --- | --- | --- |
| Crystallographic | Confidence ($t = 0.5$) | On | 2544 | 157 | 0.873 | **0.593** |
|  |  | Off | 1488 | 122 | 0.890 | 0.469 |
| | Density ($p = 0.6$) | On | 2544 | 112 | **0.938** | 0.455 |
|  |  | Off | 1488 | 112 | 0.911 | 0.442 |
| ESMFold | Confidence ($t = 0.5$) | On | 2520 | 84 | 0.789 | 0.286 |
|  |  | Off | 1488 | 81 | **0.815** | 0.286 |
| | Density ($p = 0.6$) | On | 2520 | 112 | 0.717 | **0.348** |
|  |  | Off | 1488 | 112 | 0.699 | 0.339 |

One possible reason for this is that predicted structures tend to have less ideal bond geometry than structures solved by X-ray crystallography, which has been shown to disrupt hydrogen bonds [72]. We therefore ran MolProbity [73] on both models, showing that ESMFold predicted structure has a higher MolProbity score than the deposited structure (1.69 vs 1.24), primarily due to more clashes (clash score 14.1 vs 4.65) and more poor bonds lengths (2.2% vs 0.07%). To see if subtle differences in predicted versus deposited structures result in differing water predictions, we looked for a water in the deposited structure which is within hydrogen-bonding range to at least three protein atoms, and where there were minimal deviations between the predicted and deposited atomic positions. We focused on the site defined by residues Pro66, Tyr67, Asp68, Arg98, and Gln95. The eight atoms within 4.0 Å of this water molecule differ by only 0.23 Å RMSD. In the crystal structure, at a confidence threshold of 0.5 and density of 0.6, the water molecule of interest is placed. However, in the ESMFold model, the water molecule is only placed at a density threshold of 0.6 (**Fig. 5D**).

Alternatively, the reduced performance may reflect running the predicted structure without the symmetry-mate model. To test this, we compared the deposited structure and the ESMFold model of 4RKW with and without symmetry mates. For inference with symmetry mates on the ESMFold model, we generated symmetry copies using 4RKW’s space group and unit cell. While adding symmetry mates improved the precision and recall on the crystal structure, there was little to no improvement for the ESMFold model (**Table 4**). Together, these analyses suggest that WaterFlow may be very sensitive to the local environment in placing water molecules.

Finally, we evaluated how confidence and density thresholds affected WaterFlow’s water molecule predictions near a ligand binding site. We selected a matched apo/ATP-bound structure pair of the quorum-sensing signal integrator LuxO (bound: PDB 5EP4; apo: PDB 5EP1). Both structures have resolution of 1.5 Å and the same crystallographic space group (P61). In the apo state, the deposited structure has five water molecules within the hydrogen-bonding range of binding site residues (**Fig. 5E**). Using a confidence threshold t = 0.5 recovers all five water molecules within that range, but also predicts three water molecules not observed in the deposited model. Using a density threshold ρ=0.6, three of the five water molecules within the hydrogen-bonding range are predicted, along with three water molecules not modeled in the deposited model.

In the bound structure, there are four water molecules that are within the hydrogen-bonding range with ATP, including two water molecules that serve as bridges between ATP and the protein. HOH590 links the α- phosphate oxygen (O1A, 3.0 Å) to the GLU177 side chain (OE1, 2.8 Å), and HOH538 links adenine N7 (2.6 Å) to the THR174 backbone carbonyl (2.7 Å). HOH603 is within hydrogen-bonding range to the adenine ring at N3 (2.6 Å) and HOH587 is within hydrogen-bonding range to the ribose hydroxyl (O3′, 2.8 Å). Using either inference method, t = 0.5 or ρ = 0.6, WaterFlow recovered three of the four water molecules within the hydrogen-bonding range, predicting the two bridging water molecules within 0.1Å (within experimental error) of the deposited water positions. Looking at differences between the apo and bound structure, the adenine ring in the bound structure displaces a water molecule, likely to solvate N7. WaterFlow predicts this move correctly, predicting a water molecule only 0.1 Å away from the new position in the deposited bound structure, and no water molecules predicted within 2.1 Å of the position in the apo structure. While additional analysis is needed, these results suggest that WaterFlow can pick up on critical binding site water molecules.

#### Evaluating Water Prediction on Experimental Data

Typically structure predictors are scored against deposited models, using the coordinates as ‘ground truth’ to compute precision and recall. However, these models are incomplete explanations of the underlying experimental data, with models determined by X-ray crystallography usually having around 20% residual error [36,74]. This error is thought to be in large part due to poor modeling of the protein-solvent interface [36]. This raises the question of whether precision and recall should be measured against the experimental density rather than the deposited model. A prediction scored as a false positive may correspond to a real water that was never built, and a missed water molecule may be one that should not have been modeled in the first place.

Therefore, we asked how well the predicted water molecules fit to real space electron density. To do this we evaluated the distribution of EDIAm [60] for all predicted water molecules across confidence thresholds across the 698-structure hold-out set. Lower the confidence thresholds correspond to lower median EDIAm (**Fig 6A**). At a confidence threshold of about 0.7, the median EDIAm of predicted water molecules matches the median EDIAm of deposited water molecules. A confidence threshold t = 0.9 leads to median EDIAm of predicted water molecules above that of deposited water molecules.

**Figure 6.**
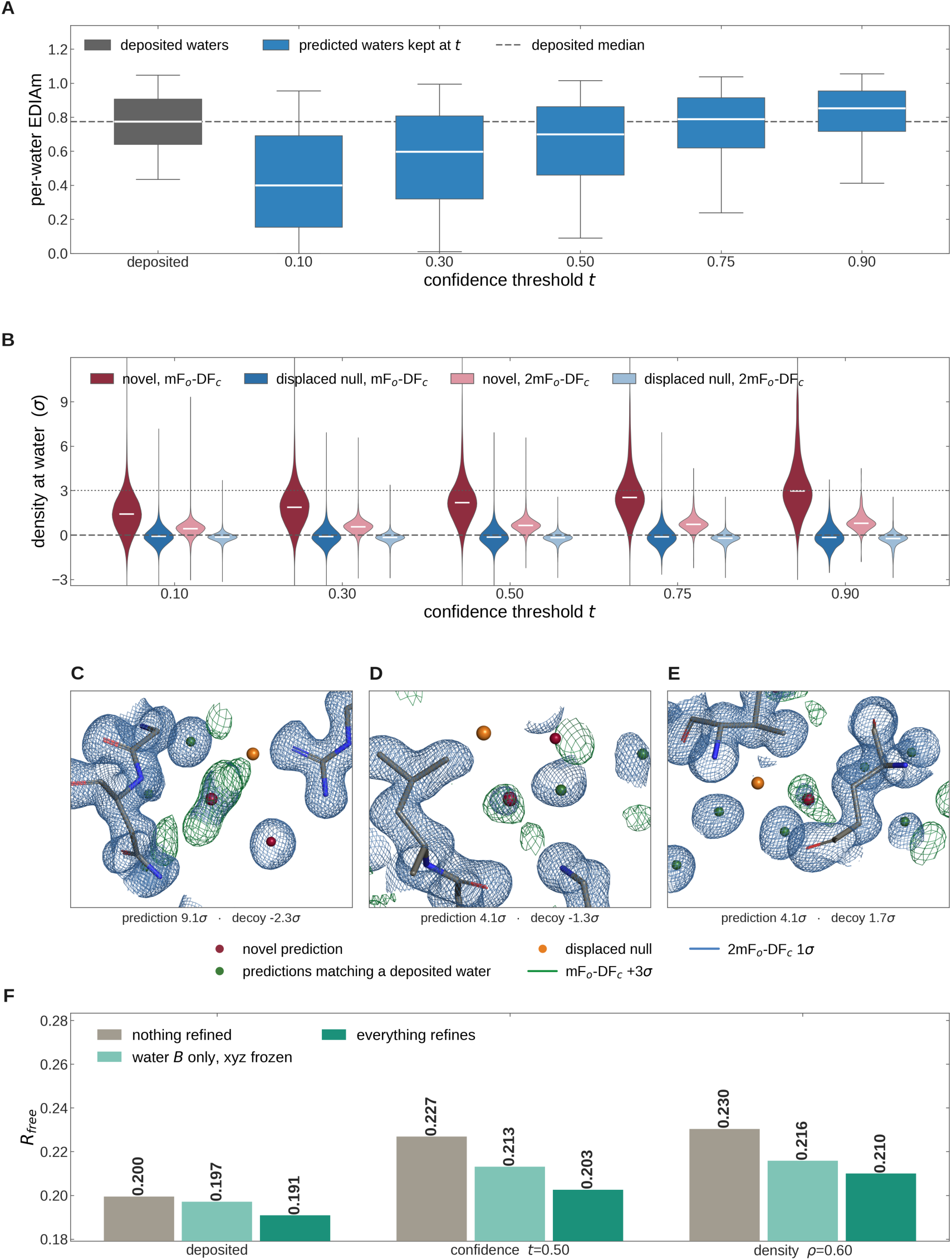
Predicted water molecules evaluated against experimental data. **A.** The distribution of per-water EDIAm for predicted water molecules across different confidence thresholds on the holdout dataset. The distribution of per-water EDIAm of the deposited water molecules (grey box). The colored boxes are the interquartile range, and the whiskers are the 5th and 95th percentile values. The deposited median EDIAm is 0.78. Median EDIAm rises from 0.40 at t = 0.10 to 0.85 at t = 0.90. **B.** Density support of novel water molecules compared to null water molecules for both the difference (mFo-DFc) and electron-density maps (2mFo- DFc) across the holdout dataset at different confidence thresholds. Red hues represent the novel water molecules, blue hues represent the null water molecules. White bars are medians, the dashed line is zero and the dotted line 3σ. The median mFo-DFc density rises from 1.41σ to 2.97σ. The fraction of novel predictions above 3σ in mFo-DFc rises from 10.7% to 49.2% as we increase the confidence threshold. The density (2mFo-DFc) median rises from 0.42σ to 0.80σ. **C-E.** Three novel water molecule predictions on PDB 4RKW at confidence threshold t = 0.5. The deposited mFo-DFc map at +3σ is green and the deposited 2mFo-DFc map at 1σ is blue, each contoured around all displayed water molecules and the decoy. The green mesh is clipped to within 2.5 Å, and the blue mesh is clipped to within 1.2 Å of any predicted water or displaced null. Novel water molecule predictions are colored crimson. Displaced null water molecules (decoys) are orange. The three sites carry 9.1σ, 4.1σ and 4.1σ against the nulls at −2.3σ, −1.3σ and +1.7σ. **F.** Refining 4RKW with predicted water molecules using a confidence threshold of 0.5 and density threshold of 0.6. We use the deposited map’s held-out reflections for three water sets, refined three ways, with coordinates fixed for the first two: i) re- scaling the bulk-solvent only, ii) only water B-factors refined, iii) full refinement of all atoms at default weights with water-picking is disabled so refinement can neither add nor delete water molecules. With nothing refined the confidence set at t = 0.5 gives R-free 0.227 against 0.200 for the deposited control, a gap of 0.027. Freeing the water B-factors narrows the gap to 0.016, and refining everything, including the protein coordinates and B-factors narrows the gap to 0.012.

We then assess the fit of novel water molecules to real space electron density, especially looking for difference density. These are water molecules predicted but not within 1.0 Å of a deposited water molecule. For each novel water molecule, we create up to eight null “decoy” water molecules randomly sampled on the surface of a 2.5 Å unit sphere around the novel water molecules, removing any nulls that overlap with any non-water heavy atom, based on van der Waals radius. Across confidence thresholds, we observed that the novel water molecules always have better real space density (2mFo-DFc), and difference density support (mFo-DFc) compared to the decoys. Similar to all water molecules, at higher confidence threshold, novel water molecules have higher density support (**Fig 6B**). At a confidence threshold of t = 0.75 the median 2mFo-DFc level reaches 3 standard deviations (3σ), the typical level where modelers would insert an atom.

Examining a select few novel water molecule predictions compared to null water molecules in 4RKW, we observe that the novel water molecules have much higher deviations of difference density (mFo-DFc) and density (2mFo-DFc) compared to null water molecules (**Fig 6C-E**).

Since real-space electron density contains phase bias [49], we then assessed our predicted water molecules against the raw structure factor amplitudes. To do this we evaluated whether WaterFlow predicted water molecules improve structure refinement. Using 4RKW, we predicted water molecules using a confidence threshold t = 0.5 (157 water molecules predicted), and using the density threshold of ρ = 0.6 (112 water molecules predicted). We assign predicted water molecules a baseline *B*-factor of 32.5 Å² (the mean value of deposited water molecules) and compute R-free. The resulting R-free values of these *unrefined* structure models with predicted water molecules is 0.227 (t = 0.5) and 0.230 (ρ = 0.6), both slightly higher than the deposited model’s R-free of 0.20. This relatively small difference indicates WaterFlow predicted water molecules fit well into the experimental data without the need for further refinement (**Fig 6F**).

We also evaluated how R-free changes when performing light refinement with the predicted water molecules. To do this, we ran phenix refinement [46] first only optimizing for the water B-factors, and then refining all coordinates, occupancies, and B-factors, including protein atoms. Water picking was turned off so no additional water molecules could be placed or removed. To make a fair comparison, we applied the same procedure to the deposited model. When optimizing water molecule B-factors, R-free improves for all models, but the changes are much larger for the two models with predicted water molecules (deposited: 0.197, t = 0.5: 0.213, ρ = 0.6: 0.216). This is probably to be expected since WaterFlow does not predict B-factors and we start by using a mean value from the deposited water molecules. When refining the complete structural model, we see a further drop in R-free for all models (deposited: 0.191, t = 0.5: 0.203, ρ = 0.6: 0.210), bringing the model with the predicted waters closer, but still having a poorer R-free. While further testing is needed, these results suggest that we may be able to use WaterFlow to model water molecules in X-ray crystallography structures.

## Discussion

Water molecules play a substantial role across biological functions, yet their impact remains understudied, in large part because modeling and predicting them has been so difficult. WaterFlow largely removes that constraint by establishing a new state of the art in predicting ordered water positions. We observe the largest gains at sub-angstrom tolerances, and we show that predicted water molecules are supported by the experimental density. This precision brings solvent prediction within reach for ligand binding analysis, seeding molecular dynamics (MD) simulations, and catalytic mechanism discovery. Treating water molecules as a predictable feature of a structure rather than a modeling afterthought opens questions across binding, resistance, and design that have been difficult to ask at scale.

We made several different choices in designing WaterFlow compared to the prior state of the art, SuperWater. The most fundamental was to use a simpler Flow Matching approach rather than Diffusion. WaterFlow’s graph structure is also considerably simpler, using only an atom-atom graph and allowing more connections between atoms. We also use ESM3 embeddings [71], which incorporate structural information, and we use a simpler GVP layer to pass messages. The simplifying choice across the model may be more appropriate for this data-limited task. While we have not exhaustively tested each alteration, we speculate that our network contains more information directly in the atom nodes, and passes that information more directly and efficiently to the water molecule nodes.

The most significant innovation of WaterFlow is the inclusion of symmetry mates, the neighboring copies of the protein generated by crystallographic symmetry operations. These copies form the crystal lattice and coordinate water molecules at the interfaces between molecules [75]. We found that adding symmetry mates to the input graph increased WaterFlow’s F1 from 0.57 to 0.63, helping most with water molecules within the hydrogen bonding range of symmetry mates. Including symmetry mates restores the coordination environment in which these water molecules are observed. Without them, a model learns to predict water positions from geometry that is systematically incomplete, and it has trouble resolving sites. We searched the literature on ligand binding, water binding, and structure prediction using both manual searches and Claude Science. Only one model includes symmetry mates or crystal contacts in its prediction context, along with discussion of this issue in PoseBuster’s dataset [24,76]. Proteins, ligands, and ions also form contacts across symmetry-related interfaces. Explicit encoding of crystallographic symmetry may therefore benefit structure prediction more broadly, particularly at interfaces and in the prediction of protein-protein interactions.

Our results also suggest what data is needed to improve water molecule predictions. Filtering water molecules based on existing metrics did not improve accuracy of the model. What did seem to drive improvements was the diversity of structures in the dataset. Additional distinct, high-quality structures are therefore the most likely route to a better performance for a water molecule prediction model.

One area where water molecule prediction is most impactful is at ligand binding sites. Water molecule displacement and solvent reorganization drive the thermodynamics of protein-ligand binding, and individual water molecules bridge protein and ligand, making accurate placement a determinant of affinity and selectivity [8,9,15]. Water molecules helping to coordinate ligand binding must be precise as hydrogen bond energies are highly sensitive to bond length, and an error of half an angstrom changes the interpretation of a network [77]. Our improvements in sub-angstrom accuracy of water molecule predictions make strong steps in the ability to predict these key connections. Using paired bound/unbound structures of LuxO [78], we demonstrate that WaterFlow has the ability to recover water molecules bridging ATP to the protein and reproducing the displacement of a water molecule by the adenine ring within experimental coordinate uncertainty. Further testing is needed to establish how well these results generalize.

Most binding site water molecules have been evaluated using physics-based solvation methods. Approaches such as 3D-RISM, GIST, and inhomogeneous solvation theory predict hydration site occupancy from a physical model of the solvent, and they return the free energy of displacing a given water [27,79,80]. WaterFlow predicts positions with a confidence score but no energetic term. Future work will test whether the two families of methods agree on which sites are occupied and whether the confidence score correlates with displacement free energies.

Across all applications, the confidence score is the main control a user has over WaterFlow’s output, and different applications will use it differently. If a user is looking for the highest precision water molecules and can tolerate low recall, say for active site water placements in QM/MM simulations [81], they can use a high confidence threshold or low density threshold. In contrast, protein designers may want to see possible water molecules across the structures and accept lower precision with better recall, and could use a lower confidence threshold or higher density threshold. Lower cutoffs result in more false positives, however our results suggest that some of the “false positives” WaterFlow predicts are ultimately supported by the experimental data. We would not recommend using a confidence threshold below 0.5, where the F1 score is still quite good.

In many of these applications, predicted structures will be used as input. However, our results indicate that WaterFlow performs worse on predicted structures compared to experimental structures. This is likely due to predicted protein structures carrying errors that WaterFlow has not been exposed to in training. WaterFlow was trained on crystallographic coordinates and has learned the local geometry characteristic of models refined, usually with geometry constraints [45,46,82]. Predicted structures carry a different distribution of coordinate error, and these small deviations have been shown to be enough to perturb hydrogen bonding patterns [72]. To overcome these errors, the confidence model likely needs recalibration for the source of the input structure, since predicted and crystallographic coordinates are not equivalent.

While we have not yet tested it, we also anticipate that WaterFlow may also have similar struggles with cryo-EM structures. While cryo-EM refinement restraints are typically the same as crystallographic restraints, which may mitigate the issue somewhat, the uncertainty in the model coordinates is larger than high-resolution X-ray crystallography structures [83,84]. Additionally, our training set includes only proteins, so ordered water molecules around nucleic acids and protein-nucleic acid complexes have not been tested and may fall outside the scope of the model [85,86]. Evaluating WaterFlow on cryo-EM and nucleic acid data will be the focus of future work.

The issues raised by WaterFlow’s performance on predicted structures identifies other issues which will need to be addressed as we push the limits of protein-bound water molecule prediction. The subtle deviations (RMSD of 0.3 Å between the crystallographic and predicted structures) are only slightly more than typical experimental coordinate uncertainty in the resolution range of our training and evaluation data (roughly 0.1 to 0.2 Å). While most PDB structural models are encoded as being static, the experimental data from which they derive capture protein and solvent that are both highly heterogeneous [87–89]. Many of the water molecules we train and evaluate against are likely partially occupied, with some of these correlating with protein fluctuations [90,91]. Models trained against static coordinates have no way to represent this [87]. Future models need to be trained with fluctuations and uncertainty in mind. This has implications far beyond water molecule prediction: *any* ligand binding will be subject to the same issues, and *any* protein prediction which is static will be limited in its utility for binding affinity prediction. Modeling protein, ligands, and solvent as a joint ensemble could recover that coupling and give access to partially occupied waters, which carry information about solvent motion and entropy that full-occupancy positions cannot represent [92,93].

However, modeling the heterogeneity inherent in proteins requires datasets that contain partially occupied water molecules alongside alternative conformers of proteins and ligands. WaterFlow is potentially a way to get this initial set of training data. WaterFlow predicted water molecules set yields R-free 0.03 higher than the deposited waters R-free; further refinement reduces the gap to 0.01 without removing or adding additional waters. This is a single demonstration and may not generalize, but it does suggest that WaterFlow could work alongside traditional refinement/model building [45–47]. Pairing it with an experimental guidance framework such as sampleworks [94], is a viable path to better data to simultaneously model water and heterogeneity. Combining model guidance with ensemble structure modeling would let solvent and macromolecule inform each other, rather than treating solvent as something added once protein and ligand are fixed.

Finally, these problems can also be viewed in light of our precision/recall “ground truth”. To determine recall and precision we used deposited water molecules as our ground truth. However, a deposited structure is itself a model of the experimental data rather than the data itself, and the water molecules modeled in it are the result of human choices [36,74]. Our ground truth is therefore imperfect, since it rests on incompletely modeled structures. A recall miss may be a water that should not have been modeled in the first place. We show predicted water molecules often well supported by electron density. The observation that novel water molecule predictions, i.e. those not near a deposited water molecule, often coincide with unexplained difference density, indicates that WaterFlow has learned at least something of the physical requirements of water molecule placements, rather than the idiosyncrasies of deposited models.

## Data and Code

Code: https://github.com/diff-use/WaterFlow

Data: 10.5281/zenodo.22046933

## Acknowledgements

This work was supported by the Radial Division at the Astera Institute (https://ror.org/050rbg919).

## Supplementary Tables and Figures

**Supplementary Table 1.** Comparing the precision and recall at 0.5 and 1 A of the generator trained on SMALL and LARGE at different water sampling ratios, where this ratio is the number of waters sampled from the flow-matching generator per residue. Evaluated on the 698 protein holdout set. GT is the ground truth number of waters.

| Model Size | Model Name | 0.5 Å |  |  |  | 1.0 Å |  |  |  |
| --- | --- | --- | --- | --- | --- | --- | --- | --- | --- |
|  |  | Precision (%) |  | Recall (%) |  | Precision (%) |  | Recall (%) |  |
|  |  | Water Ratio |  |  |  |  |  |  |  |
|  |  | GT | 5 | GT | 5 | GT | 5 | GT | 5 |
| SMALL | SuperWater | 7.8 | 2.6 | 7.8 | 23.9 | 18.6 | 7.1 | 18.3 | 57.3 |
|  | WaterFlow | 33.3 | 29.6 | 24.2 | 72.9 | 45.2 | 40.0 | 31.9 | 85.5 |
|  | WF (no contacts) | 24.8 | 22.0 | 19.3 | 60.1 | 34.9 | 30.7 | 26.5 | 74.9 |
| LARGE | SuperWater | 9.4 | 2.0 | 9.4 | 19.4 | 22.2 | 6.3 | 21.8 | 54.2 |
|  | WaterFlow | **34.0** | **31.2** | **24.5** | **73.2** | **46.1** | **41.8** | **32.4** | **86.1** |
|  | WF (no contacts) | 29.7 | 27.1 | 21.7 | 60.3 | 40.2 | 36.3 | 28.7 | 74.1 |

**Supplementary Table 2-4** Information on the datasets used for consensus water analysis (PDB ID, number of water, resolution, R-free, etc.): https://zenodo.org/22046933

**Supplementary Table 5:** Consensus water clustering statistics. The fraction of the consensus waters in each dataset from the best clustering result is reported along with its range across the hyperparameter grid search (see **Methods**). Within each dataset, structures are binned into 10 equal-count bins by the number of water molecules modeled, and the average P/R (precision/recall) in the lowest and highest bins are reported.

| dataset | Best clustering hyperparameters<br>(min_cluster_size,<br>min_sample_size) | Best consensus water % | Range of consensus water % | P/R lowest num_water bin | P/R highest num_water bin | P/R Pareto knee |
| --- | --- | --- | --- | --- | --- | --- |
| HEWL | 5, 5 | 69.85 | 63.22 - 70.35 | 0.94 / 0.36 | 0.47 / 0.85 | 0.97 / 0.78 |
| EAP | 20, 20 | 86.5 | 83.58 - 86.5 | 0.93 / 0.47 | 0.70 / 0.89 | 0.92 / 0.96 |
| CA | 15, 5 | 78.55 | 77.83 - 79.06 | 0.92 / 0.36 | 0.68 / 0.84 | 0.88 / 0.86 |

| dataset | Best clustering hyperparameters<br>(min_cluster_size,<br>min_sample_size) | Best consensus water % | Range of consensus water % | P/R lowest num_water bin | P/R highest num_water bin | P/R Pareto knee |
| --- | --- | --- | --- | --- | --- | --- |
| HEWL | 5, 5 | 62.28 | 48.93 - 62.28 | 0.92 / 0.39 | 0.39 / 0.80 | 0.91 / 0.81 |
| EAP | 20, 20 | 83.86 | 81.32 - 83.86 | 0.87 / 0.45 | 0.66 / 0.86 | 0.91 / 0.94 |
| CA | 15, 5 | 70.44 | 70.23 - 70.82 | 0.82 / 0.36 | 0.59 / 0.81 | 0.87 / 0.86 |

**Supplementary Table 6:** Distribution of the resolution and R-free of structures above versus below the median F1 score (HEWL: 0.65, EAP: 0.84, CA: 0.69) against the consensus set of water molecules for each dataset. This table corresponds to Table 2 in the main text, except that a distance cutoff of 0.5 Å is used for clustering the consensus waters and for computing the F1 scores.

| metric | dataset | above |  | below |  | n_above | n_below | r | p |
| --- | --- | --- | --- | --- | --- | --- | --- | --- | --- |
|  |  | median | iqr | median | iqr |  |  |  |  |
| resolution (Å) | HEWL | 1.55 | (1.45, 1.63) | 1.6 | (1.35, 1.8) | 32 | 33 | -0.08 | 0.58 |
|  | EAP | 1.16 | (1.08, 1.26) | 1.43 | (1.18, 1.62) | 454 | 453 | -0.53 | 2.26×10<sup>-43</sup> |
|  | CA | 1.35 | (1.2, 1.5) | 1.5 | (1.23, 1.7) | 468 | 469 | -0.28 | 1.85×10<sup>-13</sup> |
| deposited R-free% | HEWL | 18.57 | (17.3, 20.12) | 20.68 | (18.86, 22.14) | 32 | 33 | -0.41 | 4.69×10<sup>-3</sup> |
|  | EAP | 15.81 | (14.84, 17.33) | 17.37 | (15.37, 19.23) | 454 | 453 | -0.3 | 1.91×10<sup>-15</sup> |
|  | CA | 17.02 | (15.47, 18.24) | 17.82 | (15.9, 19.61) | 468 | 469 | -0.2 | 1.55×10<sup>-7</sup> |

**Supplementary Table 7:** Distribution of the B-factor z-score and the EDIA of consensus versus non-consensus water molecules in each dataset. This table corresponds to Table 3 in the main text, except that a distance cutoff of 0.5 Å is used for clustering the consensus waters.

| metric | dataset | consensus |  | nonconsensus |  | n_consensus | n_nonconsensus | r | CI |
| --- | --- | --- | --- | --- | --- | --- | --- | --- | --- |
|  |  | median | iqr | median | iqr |  |  |  |  |
| B-factor z-score | HEWL | -0.44 | (-0.95, 0.15) | 0.6 | (-0.02, 1.17) | 3605 | 2183 | -0.57 | (-0.61, -0.54) |
|  | EAP | -0.21 | (-0.89, 0.41) | 0.89 | (0.34, 1.44) | 191272 | 36809 | -0.62 | (-0.63, -0.6) |
|  | CA | -0.33 | (-0.88, 0.26) | 0.64 | (0.06, 1.23) | 151731 | 63670 | -0.56 | (-0.57, -0.55) |
| EDIA | HEWL | 0.8 | (0.61, 0.93) | 0.6 | (0.4, 0.77) | 3605 | 2183 | 0.42 | (0.36, 0.48) |
|  | EAP | 0.89 | (0.76, 0.99) | 0.6 | (0.45, 0.73) | 191272 | 36809 | 0.72 | (0.71, 0.73) |
|  | CA | 0.83 | (0.7, 0.94) | 0.61 | (0.46, 0.74) | 151730 | 63670 | 0.57 | (0.56, 0.58) |

**Supplementary Table 8:**
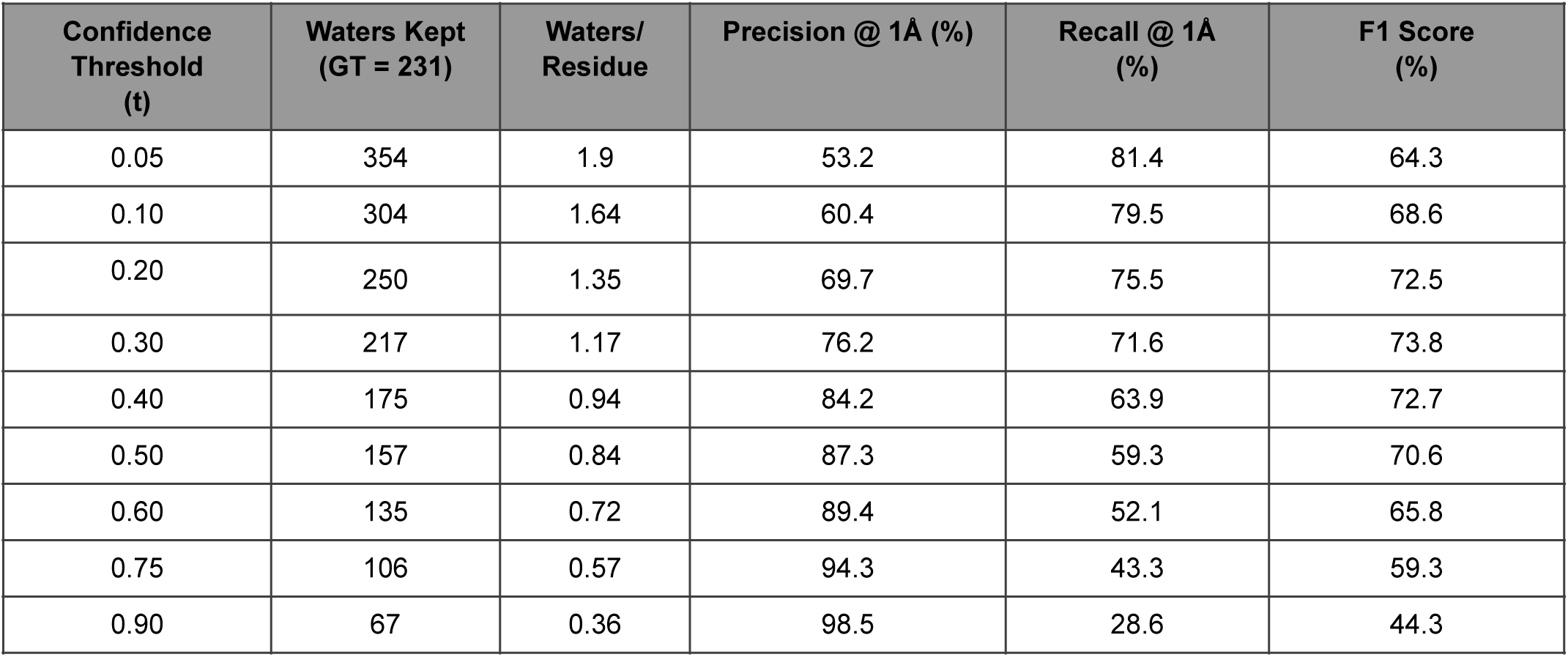
Prediction statistics with confidence thresholding for PDB 4RKW, inference on symmetry-mates model checkpoint.

**Supplementary Table 9:** Prediction statistics with confidence thresholding for PDB 4RKW, inference on symmetry-mates-off model checkpoint.

| Confidence Threshold (t) | Waters Kept (GT = 231) | Waters/Residue | Precision @ 1Å (%) | Recall @ 1Å (%) | F1 Score (%) |
| --- | --- | --- | --- | --- | --- |
| 0.05 | 336 | 1.81 | 47.6 | 69.3 | 56.4 |
| 0.10 | 275 | 1.48 | 55.7 | 66.2 | 60.5 |
| 0.20 | 205 | 1.1 | 66.6 | 59.2 | 62.7 |
| 0.30 | 164 | 0.88 | 77.6 | 55.0 | **64.4** |
| 0.40 | 138 | 0.74 | 84.6 | 50.6 | 63.4 |
| 0.50 | 122 | 0.65 | 89.0 | 46.9 | 61.4 |
| 0.60 | 104 | 0.56 | 91.6 | 41.1 | 56.8 |
| 0.75 | 76 | 0.41 | 93.0 | 30.6 | 46.0 |
| 0.90 | 47 | 0.25 | 96.5 | 19.8 | 32.8 |

**Supplementary Table 10:**
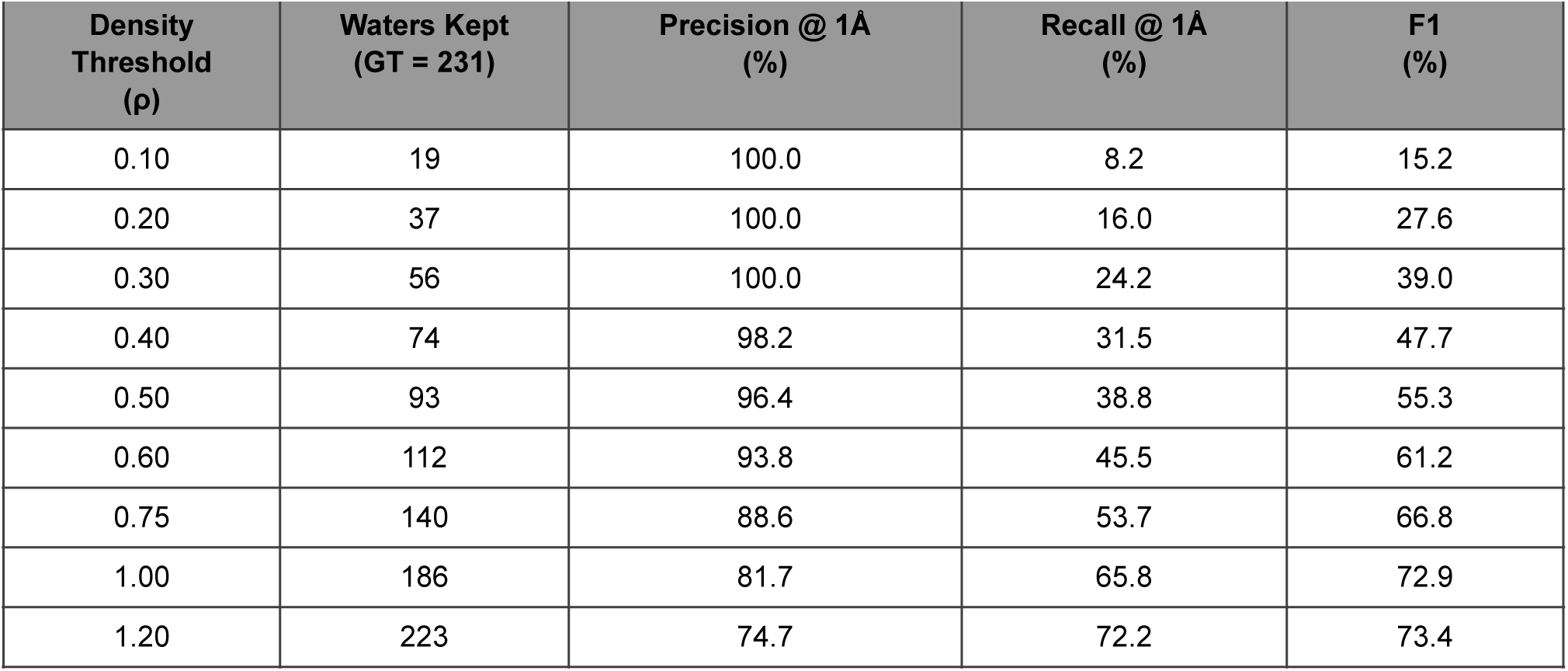
Prediction statistics with density thresholding for PDB 4RKW, inference on symmetry-mates model checkpoint.

**Supplementary Table 11:**
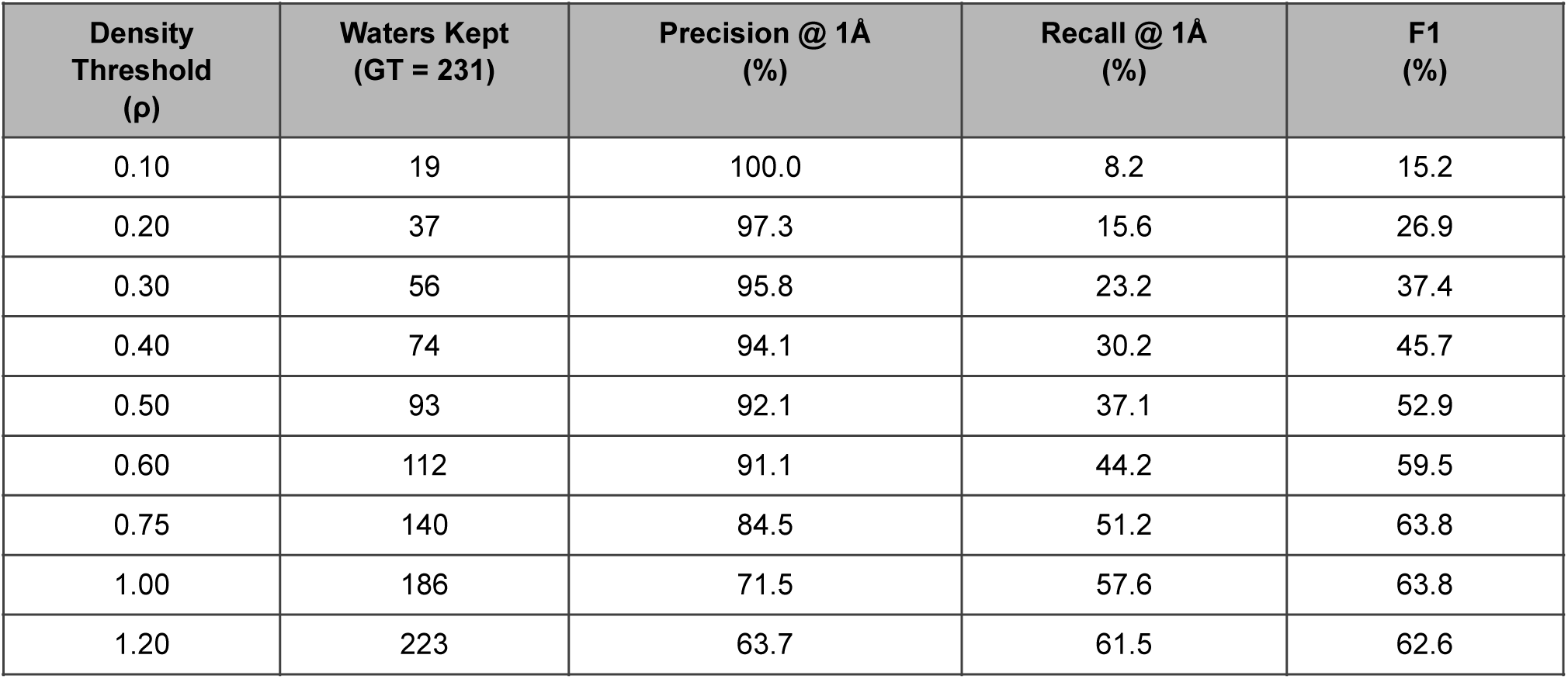
Prediction statistics with density thresholding for PDB 4RKW, inference on symmetry-mates-off model checkpoint. Predictions for ST8-11 are computed against waters that pass quality filters, with 9 waters failing the B-factor z-score ≤ 2.0 filter, and 6 waters failing the EDIAm ≤ 0.4 filter. 5 waters fail both rules. 231 out of 242 waters remain.

**Figure S1.**
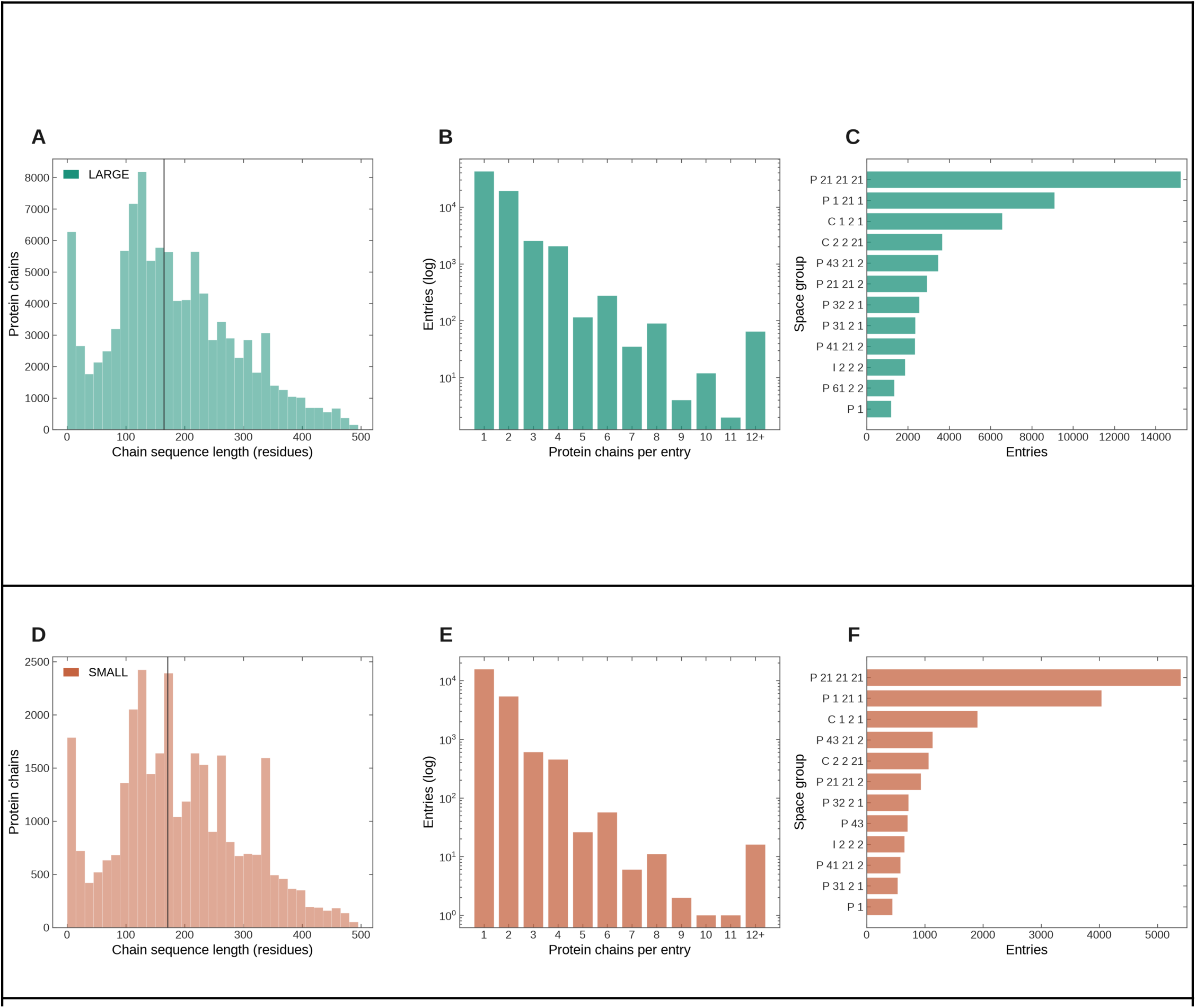
Sequence, assembly, and crystal-form composition of the two training datasets. LARGE (teal, A–C) and SMALL (terracotta, D–F). The datasets are clustered into families by sequence (MMseqs2, ≥ 30% identity, ≥ 80% coverage) and structural (Foldseek multimer, TM ≥ 0.5) similarity before splitting, giving 15,573 clusters over LARGE and 5,424 over SMALL (Methods). **A, D. Chain Sequence Length.** One count per protein chain over all chains (LARGE 101,574 chains, SMALL 31,038), binned every 15 residues. The median chain length (black bar) is 165 residues (interquartile range 107–247) for LARGE and 171 residues (116–259) for SMALL. The two datasets have near-identical chain length composition. **B, E. Protein chains per entry.** Number of protein chains per deposited entry, counts on a log axis, with 12 or more pooled into the final bin. Both datasets are dominated by monomeric and dimeric entries, with higher-order assemblies falling off by roughly an order of magnitude per added chain. **C, F. Space groups.** The twelve most frequent space groups by entry count, as horizontal bars in descending order. P 2₁ 2₁ 2₁ is the most common in both datasets by a wide margin, followed by P 1 2₁ 1 and C 1 2 1, with near-identical ordering across LARGE and SMALL.

**Figure S2.**
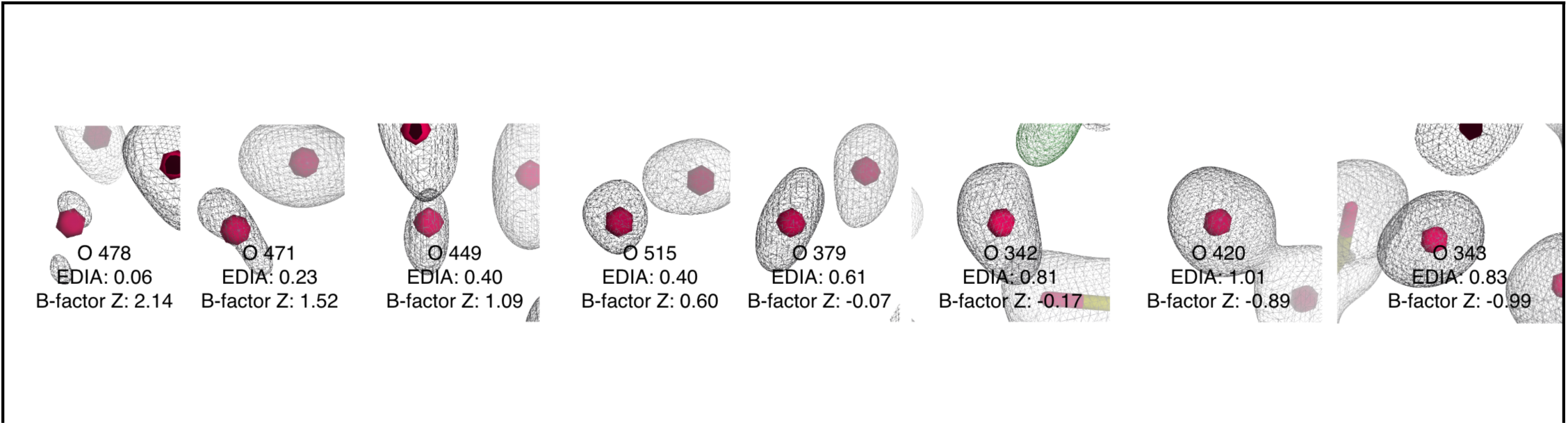
Examples of water molecules with varying EDIA and B-factor z-score shown with the 2Fo-Fc electron density map.

**Figure S3.**
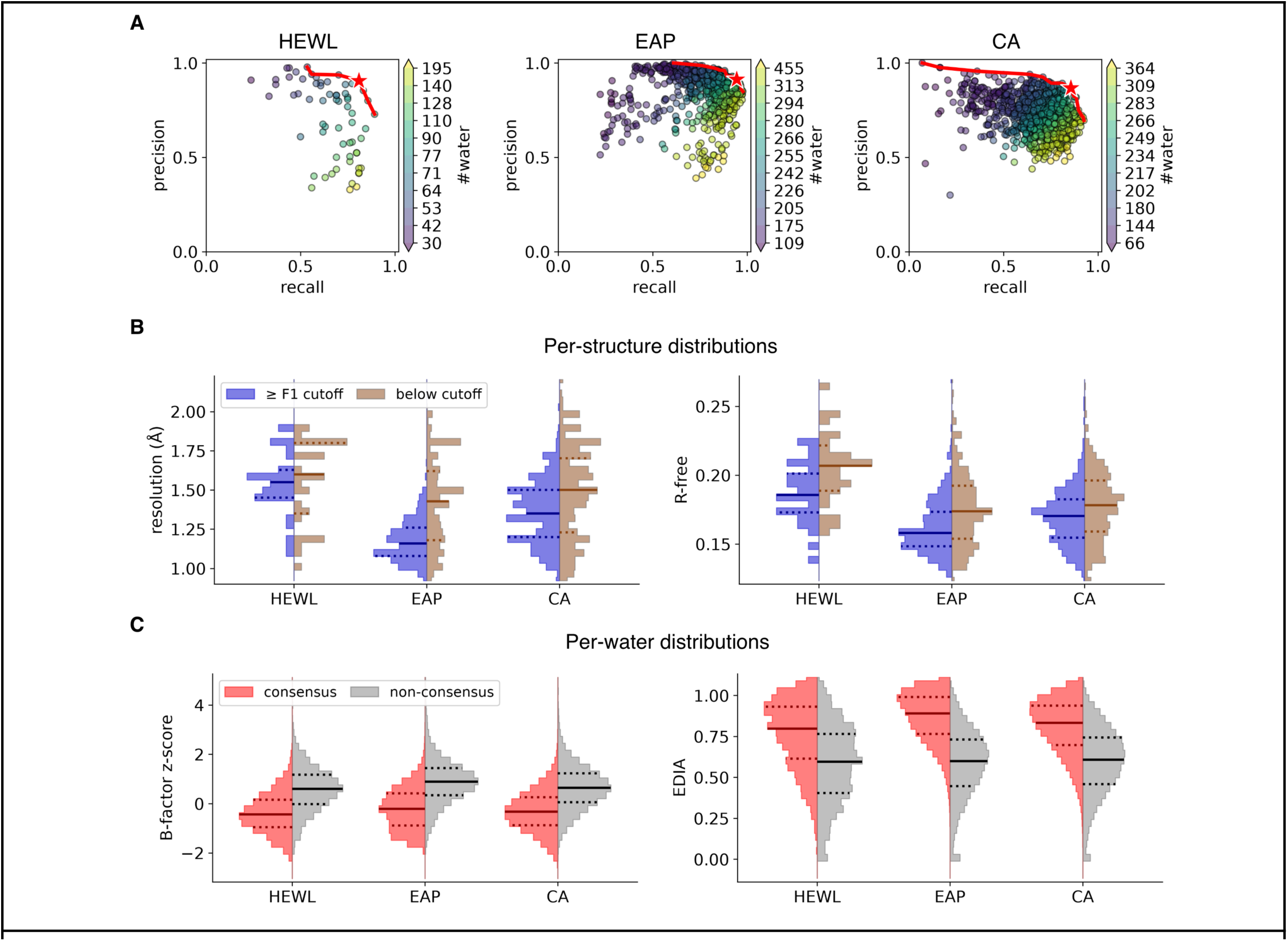
Consensus water analysis corresponding to Fig. 2B-D with a cutoff distance of 0.5 Å is used. Clustering statistics are shown in Supplementary Table 5. **A.** The Pareto knee in panel B has precision/recall/F1 of 0.91/0.81/0.86 for HEWL, 0.91/0.94/0.93 for EAP, and 0.87/0.86/0.86 for CA. **B.** For the per-structure metric, the better half of structures (purple) also has higher resolution (median 1.55 vs 1.60 Å for HEWL, 1.16 vs 1.43 Å for EAP, 1.35 vs 1.50 Å for CA) and lower R-free (median 18.6% vs 20.7%, 15.8% vs 17.4%, 17.0% vs 17.8%). **C.** Consensus water molecules (red) have lower B-factor z-score (median -0.44 vs 0.60 for HEWL, -0.21 vs 0.89 for EAP, -0.33 vs 0.64 for CA) and higher EDIA (median 0.80 vs 0.60 for HEWL, 0.89 vs 0.60 for EAP, 0.83 vs 0.60 for CA) than the non-consensus water molecules.

**Figure S4.**
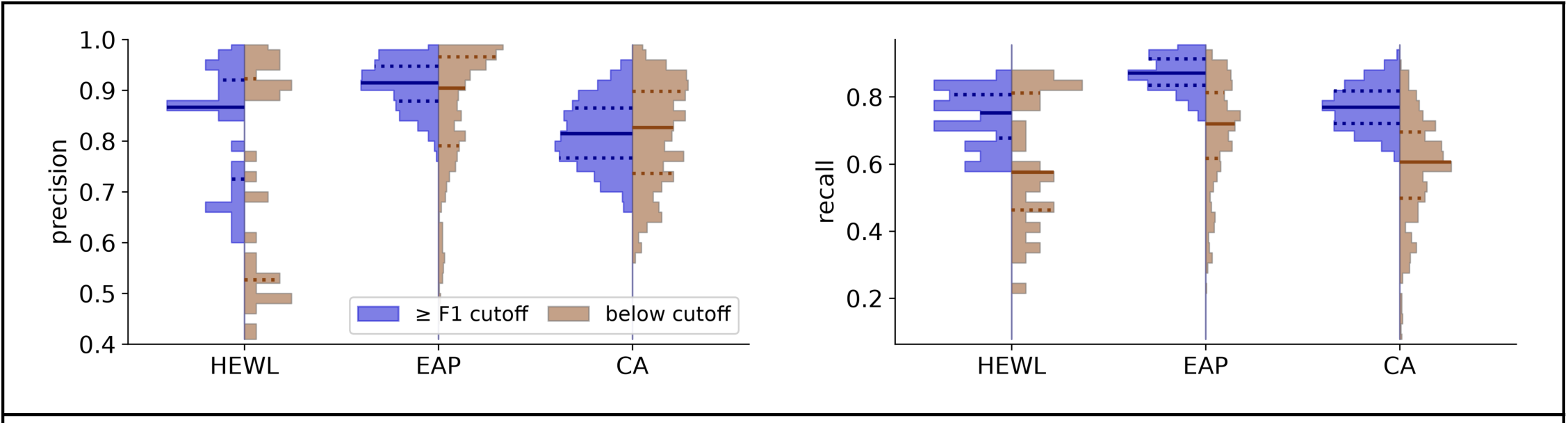
Distribution of the precision and recall of each dataset split by median F1 score against consensus water (distance cutoff = 1.0 Å). Each distribution’s median value and IQR are marked by solid and dotted lines respectively.

**Figure S5.**
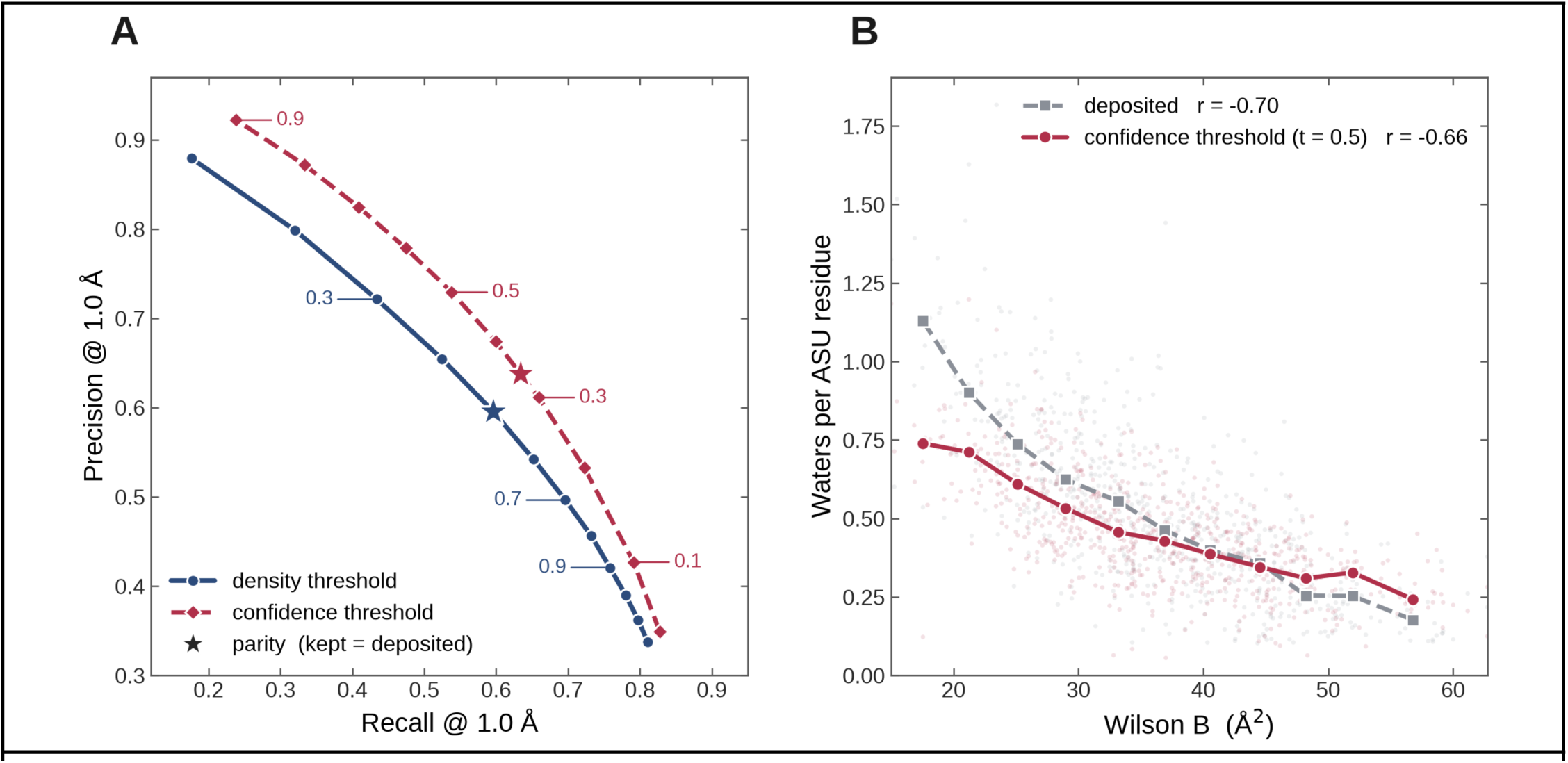
**A.** Precision/Recall at 1Å. Each curve is a method swept across its thresholds. Labelled markers give the threshold value (density in navy, confidence in crimson). The confidence curve lies above the density curve throughout, holding roughly 0.05–0.08 higher precision at matched recall. The stars mark parity on each curve, which is the operating point at which total water molecules predicted equals total deposited. Confidence thresholding retains higher precision even at parity. Maximum precision and recall for the density curve are 0.879 and 0.811 at ρ = 0.1 and ρ = 1..2 respectively, and a best F1 score of 0.596, compared to the confidence thresholding metrics reported in Fig 1B. **B. Water molecules per ASU residue against Wilson B.** Water molecule count normalized per asymmetric-unit residue versus the structure’s Wilson B factor (global crystallographic disorder). Faint points are individual structures, joined markers are medians within thirteen Wilson-B bins spanning the 1st–99th percentile (≥ 12 structures per bin). Grey squares are the deposited water molecule count, correlating strongly with the Wilson B-factor (Pearson r = −0.70). Crimson circles are the confidence threshold predictions at t = 0.50, which also correlate strongly with the Wilson B- factor (Pearson r = −0.66).

**Figure S6.**
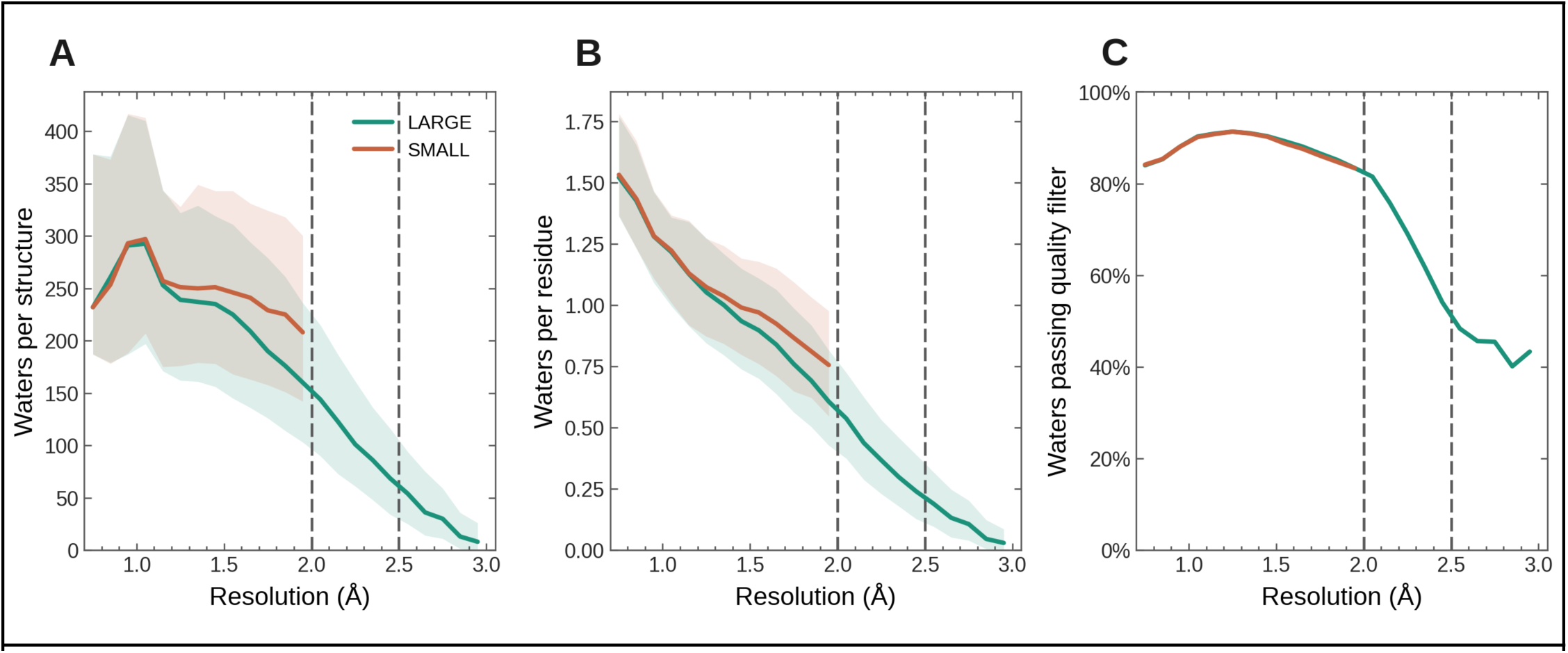
LARGE (teal, resolution ≤ 3.0 Å) and SMALL (terracotta, ≤ 2.0 Å). Curves are medians within 0.1 Å resolution bins. **A. Water molecules per structure as a function of resolution.** Median count of deposited water molecules per structure. Near 1.0 Å resolution we observe close to ∼290-300 water molecules. Post 1.0 Å we see a sharp fall, the LARGE median drops from ∼265 water molecules below 1.2 Å to ∼29 in the 2.5–3.0 Å bin. SMALL sits slightly above LARGE (∼212 vs ∼161 in the 1.8–2.0 Å bin) where they overlap, consistent with its higher-resolution, better-modeled structures carrying more modelled water molecules. **B. Water molecules per residue as a function of resolution.** We evaluate the same deposited water molecule count normalized by the number of residues in the asymmetric unit of each structure. The LARGE median falls monotonically from ∼1.18 water molecules per residue above 1.2 Å to ∼0.10 by 2.5–3.0 Å. The SMALL median falls from ∼1.20 below 1.2 Å to ∼0.77 in the 1.8–2.0 Å bin, and sits above LARGE in every overlapping bin (∼0.77 vs ∼0.61 at 1.8–2.0 Å). **C. Fraction of deposited water molecules passing quality filters as a function of resolution.** Share of deposited water molecules clearing both filters (EDIAm ≥ 0.4 and B-factor z ≤ 2.0) on a per-water rate. Around 90% of deposited water molecules are kept to ∼1.5 Å resolution and 84% to 2.0 Å, then falling to roughly 45% by 2.5–3.0 Å. LARGE and SMALL are indistinguishable where they overlap.

**Figure S7.**
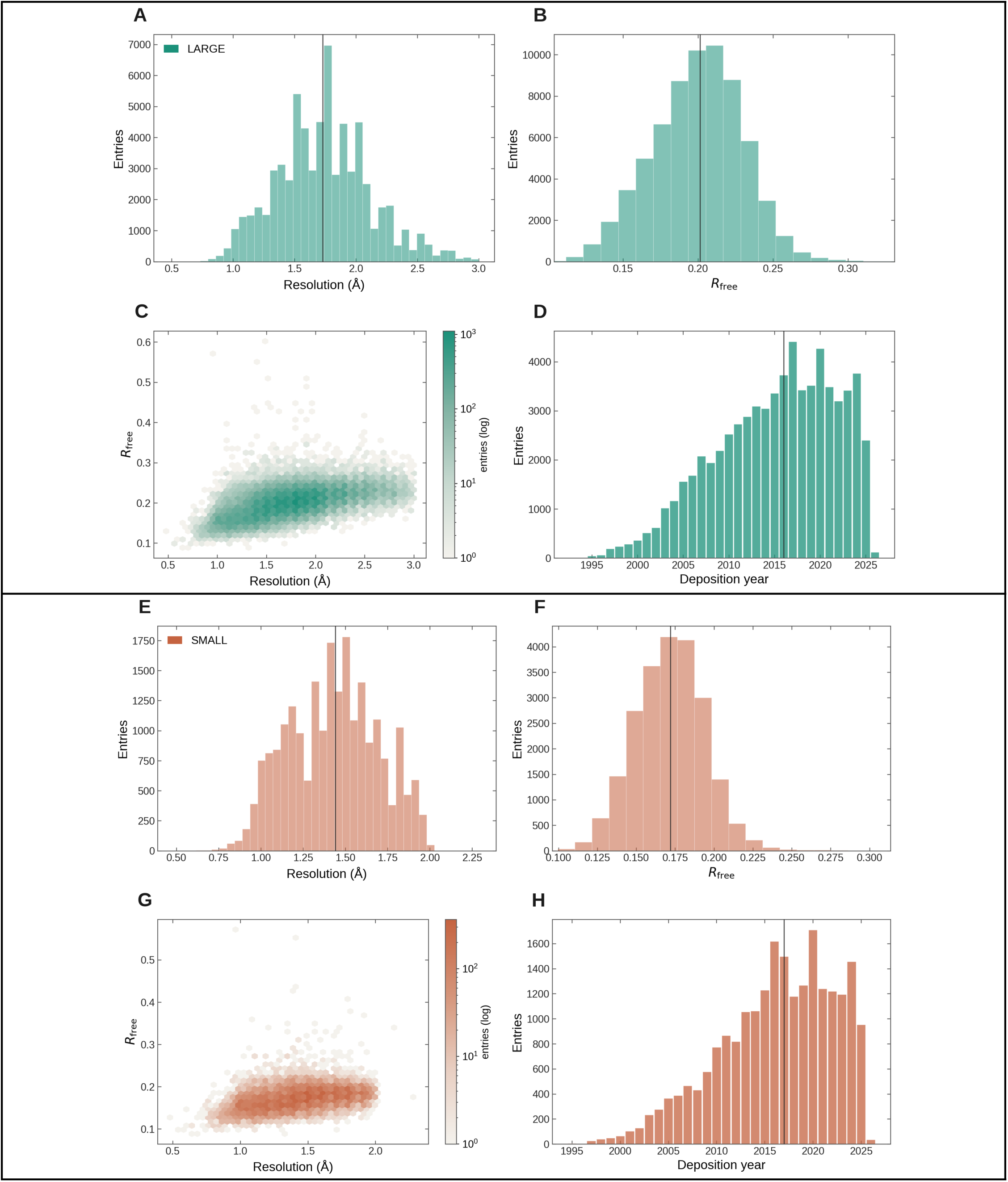
Crystallographic quality of the LARGE and SMALL datasets. Both datasets are obtained from PDB-REDO re-refined structures. LARGE (67,272 entries, resolution ≤ 3.0 Å) is in teal, and SMALL (22,318 entries, ≤ 2.3 Å) is in terracotta. Panels A–D describe LARGE and E–H describe SMALL. Every quantity is per deposited entry; the vertical black rule in the histograms marks the median. **A, E. Resolution.** Distribution of crystallographic resolution (40 bins). LARGE spans 0.48–3.00 Å with a median (black line) of 1.73 Å. SMALL spans 0.48–2.30 Å with a median of 1.44 Å. **B, F. R-free.** Distribution of R-free values on both datasets. LARGE has a median R-free of 0.201 with 90% of entries below 0.236 and 99% below 0.268. SMALL SMALL has a median of 0.172, with 90% of entries below 0.199 and 99% below 0.227. **C, G. Resolution against deposited R-free.** Hexbin density color is entry count on a log scale. The two metrics are positively associated but have significant spread. **D, H. Deposition Year.** Entries per year of deposition. Both datasets weight towards recent years (median 2016 for LARGE and median 2017 for SMALL), tracking the growth of high-resolution depositions.

**Figure S8.**
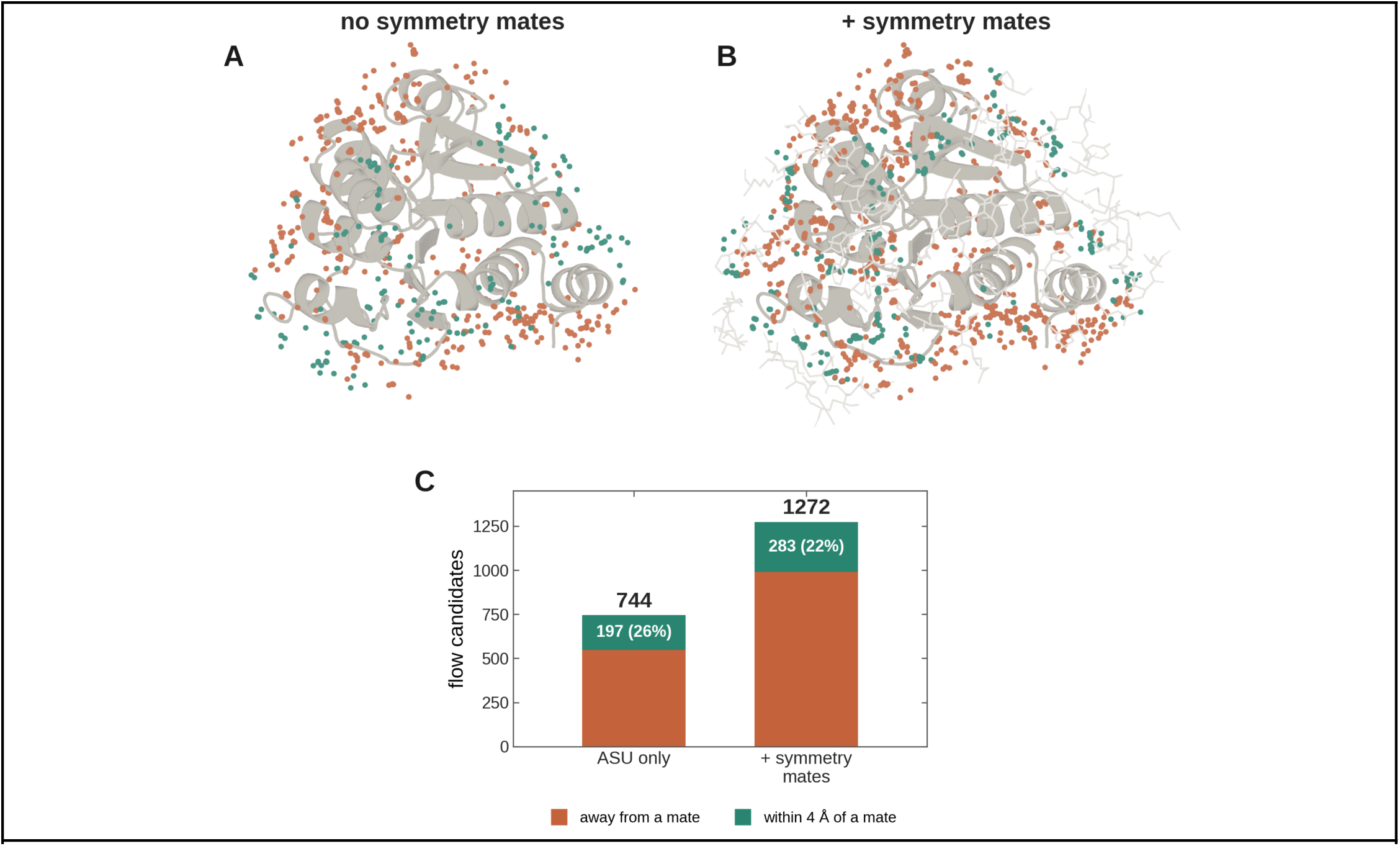
Symmetry mates enlarge and redirect the flow-matching generator’s candidate cloud. The flow-matching generator is run twice on 4RKW at the same water to residue ratio (4.0), differing only in whether symmetry mates exist in the graph. **A.** Candidate water molecules with ASU-only graph (186 residues). Out of 744 total candidates, 197 (26%) land within 4Å of a symmetry mate. **B.** Candidate water molecules with symmetry mates included in the graph (318 residues). Symmetry-mate ghosts are shown as sticks. Candidates are coloured by proximity to a mate. Terracotta: > 4 Å distance from a symmetry mate, teal ≤ 4 Å of symmetry mate. Out of 1272 candidates, 283 (22%) lie within 4Å of a symmetry mate. **C.** Candidate water molecules count per run. Because the water molecule-to-residue ratio scales with the number of graph residues, the mate’s graph samples 528 more candidates, of which 283 (22 %) fall within 4 Å of a symmetry mate. These are positions that the ASU-only graph has no context to propose and hence the added symmetry mate residues aid in increasing recall.

**Figure S9.**
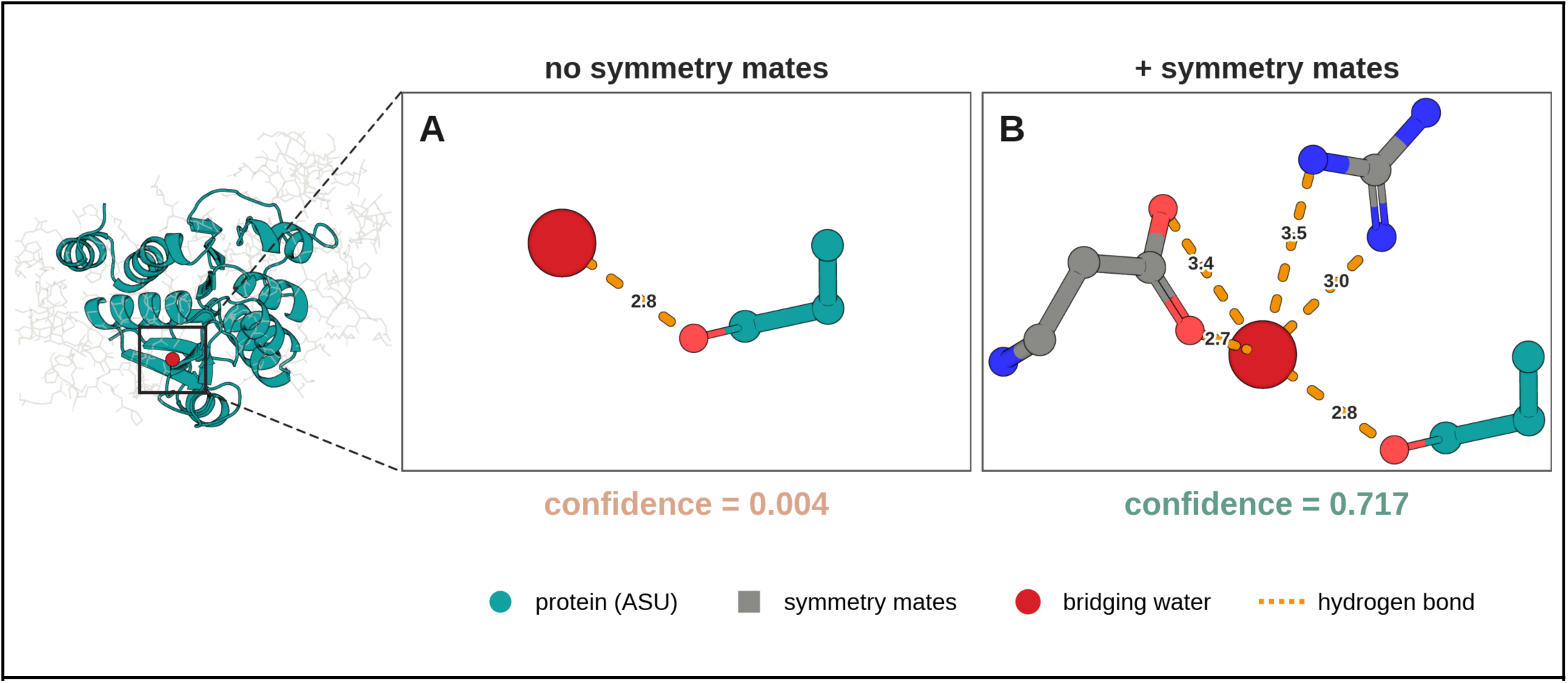
Symmetry mates lead to higher confidence scores for bridging waters. A single deposited water molecule in a symmetry-mate contact of 4RKW (DJ-1/PARK7, 1.50 Å, 186 ASU residues), scored by the confidence head in two graphs that differ only in whether symmetry mates are present. Left, the DJ-1 fold (teal) within its crystal environment (grey symmetry-mate ghosts), with the site boxed and inset to the zoom panels. Teal is the protein ASU. Grey is a symmetry mate. Red is the bridging water. Orange dotted lines are potential hydrogen bonds (≤ 3.4 Å) labelled in Å. Coordinating residues are trimmed to their water-facing portion. This is a single-site counterpart of the dataset-level statistic in the main text Fig. 4. **A.** ASU only: the water molecule retains its single ASU potential hydrogen bond (2.8 Å) and has a confidence score of 0.004. **B.** The symmetry mates contribute four additional potential hydrogen bonds (2.7 – 3.5 Å) and the same water molecule has a confidence score of 0.717.

