## Supplementary material for "WaterFlow: Prediction of Ordered Water Molecule Positions on Protein Structures": Methods

### WaterFlow Supplementary Methods

Overview of the method

WaterFlow places ordered water molecules on a protein structure in two stages: a flow-matching generator proposes candidate positions, and a confidence model selects among them. Both stages use geometric vector perceptron (GVP) graph networks [5] and are equivariant to rigid motions of the input. Inference has four steps.

1. The protein, retained ligands, and crystallographic neighbors that contact the asymmetric unit (ASU) form an all-atom graph. Protein atoms carry an ESM3 residue embedding fused with a one-hot element identity; ligand atoms have a zero residue embedding.
2. The generator samples candidate water positions. It draws $r = 8$ candidates per residue represented in the graph.
3. The confidence model assigns each candidate a score in $[0, 1]$.
4. Candidates below a score threshold are removed. A two-pass rule at 1.52 Å then merges the survivors into discrete waters.

Sweeping the threshold in step 4 traces a precision–recall curve and controls the number of returned waters. Section 7.3 also evaluates a density thresholding alternative.

#### 1 Protein structure selection and pre-processing

##### 1.1 PDB-REDO subset

We chose to work with protein structures from PDB-REDO [20, 6]. PDB-REDO applies consistent refinement and rebuilding protocols to solve structures from X-ray diffraction data. It takes advantage of hydrogen-bonding information which should be important to water binding to protein. PDB-REDO contains more than 100,000 structures. We selected protein structures by requiring 50-500 amino acid residues in at least one polypeptide chain.

To test the impact of data quantity on model quality, we constructed two subsets from the initial query, based on the X-ray resolution and the model quality reflected in $R_{\text{free}}$. The Large data comprises all structures from our initial subset with resolution $\leq 3.0$ Å and $R_{\text{free}} \leq 0.25$, yielding 67,272 entries. The Small data is further restricted to the 22,318 structures with resolution $\leq 2.0$ Å and $R_{\text{free}} \leq 0.20$. All ligands, ions, and co-factors are retained in the selected entries to provide WaterFlow with the same context as is present in the original data.

##### 1.2 Dataset partitioning

We partitioned structures using a combination of sequence and structure clustering; each cluster is then assigned to training, testing, or validation data. Clustering is done by constructing a graph of connections between proteins based on similarity, and identifying connected subgraphs.

We use MMseqs2 [18] to compare all protein sequences. A sequence edge connects two entries when a one-to-one chain assignment has at least 30% sequence identity and 80% coverage per matched chain and covers at least 50% of the residues in each complex. The chain assignment maximizes total alignment score, so complex coverage is residue-weighted.

We apply Foldseek multimer alignment [21] separately to each sequence-connected component. We include an edge in the graph if the minimum TM-score between two protein structures is at least 0.5. (TM-scores are not symmetrical, so we compute the score using each protein as the target). We require 80% coverage requirement.

Finally we keep only those edges in the graph that have both a sequence and structural similarity edge. Final clusters are the connected components of the full graph. The LARGE selection contains 67,272 structures in 15,573 clusters with a mean size of 4.32 and 68.6% singletons. The SMALL selection contains 22,318 structures in 5,424 clusters with a mean size of 4.11 and 68.3% singletons.

Clusters are assigned to achieve an approximate ratio 95/2.5/2.5 train/validation/test by structure count. The validation and test sets are built with whole clusters containing 50–100 structures to balance diversity and the ability to analyze whole clusters to determine consensus between different related proteins. Clusters larger than 100 structures remain in training. LARGE's final split is 63,901 training, 1,688 validation, and 1,683 test structures. The corresponding SMALL counts are 21,270, 506, and 542.

To ensure that test and validation sets can be used with any model, any SMALL cluster that spans more than one LARGE split is moved to the SMALL training set. This can happen because of slight variances in the MMseqs2 algorithm. It affects only 33 structures total.

The union of the SMALL validation and test assignments defines a 1,048-structure high resolution set that we use for hyperparameter selction and validation. WaterFlow graphs are available for 1,044 structures, and SuperWater clouds are available for 1,041. The missing structures fail the corresponding pipeline's processing criteria.

For the model comparisons reported below we use a holdout that no model was trained or selected on. Starting from the LARGE test assignment we remove every structure used to train or select any model, leaving 698 structures.

##### 1.3 Additional filtering

We retained water molecules from PDB-REDO structures if they met three conditions:

1. The water oxygen falls within 5.0 Å of a protein heavy atom,
2. it has an EDIAm $\geq 0.4$ [14], and
3. it has a B-factor z-score $\leq 2.0$, relative to the waters in that structure.

When we include symmetry-mate atoms (§1.5) in the structure, condition 1 applies to those atoms as well; i.e., a water is retained if its oxygen is within 5.0 Å of any *polypeptide* atom included in the atomic graph as defined in section §1.6. This explicitly *does not* include ligands: a water molecule that is within 5.0 Å of a ligand atom but not within that distance of any protein atom is not included.

We then remove whole structures that don't meet the following quality criteria:

1. At most, 5% of water molecules may lie within 2.0 Å of a protein atom (including symmetry mates), since this may indicate poorly-modeled solvent or a coordinate-frame error.
2. At minimum, the structure must contain 0.1 ordered water molecules per amino acid residue in the polypeptide chain(s), which can indicate problems with the X-ray data or model refinement [22].
3. Multi-chain entries must contain at least one inter-chain contact, defined as having one atom in the first chain within 4.0 Å of an atom in the second chain.

The per-water filtering removes a median of 23 (25) waters per structure in the LARGE (SMALL) dataset. The whole-structure criteria are applied after the train/validation/test split, so their effect is reported there against each split rather than against the full selection.

The water and structure filters applied reduce the LARGE training list from 63,901 to 60,118 structures and the SMALL training list from 21,270 to 21,211. Both the generator and confidence model of each dataset were trained on the full cached training list. Validation and test structures are used as assigned in §1.2.

##### 1.4 Parsing

Structures are parsed with Biotite [8]. If more than one model is present, we select the first model only. When there is more than one alternate conformer, we retain the highest-occupancy one. We retain modified residues, such as selenomethionine. All hydrogen atoms are removed. B-factors and EDIAm scores are joined in from PDB-REDO metadata files; we drop structures without accompanying metadata files.

##### 1.5 Generating crystal contacts

The structure reported we use from a PDB-REDO entry is generally the *asymmetric unit*. The complete structure of the protein crystal includes the *space group*, which describes transformations of the asymmetric unit required to complete the unit cell, and the dimensions of the unit cell itself which define the crystallographic lattice. Crystallographic neighbors are generated with the PyMOL[17] command `symexp` using the deposited space group and unit cell. We keep all protein and ligand atoms that are within 8.0 Å of any atom in the asymmetric unit. We do not retain water molecules that are generated as a result of symmetry operations. Atoms within 0.3 Å of an ASU atom, a target water, or another already symmetry-transformed copy of the ASU are removed. Symmetry-generated ligands and residues are retained whole.

##### 1.6 Graph construction

###### 1.6.1 Edge construction

Surviving structures are converted to heterogeneous graphs, consisting of context nodes (protein and ligand atoms) and water nodes, expressed in a frame centered on the ASU protein-atom centroid. By the translation-equivariance of the architecture (§2) this normalisation does not affect the predicted water positions. They translate with the frame and are mapped back to the input crystal frame at output (§9). We hence use a centroid basis as a coordinate convention rather than a modeling choice.

Protein–protein edges are built once during pre-processing and cached. A directed edge is added between every protein atom and every other protein atom within 8.0 Å, retaining at most 256 neighbors per atom. We then add the reverse of every edge and drop duplicates. We add edges ending at water nodes in the same fashion, but we do not add the reverse edges. Edges including water nodes are not cached, as their positions change during flow integration.

###### 1.6.2 Node and edge features

Each node stores a 16-dimensional one-hot vector over the element vocabulary, and a residue index if the atom belongs to a polymer residue. Each node also stores boolean flags for symmetry mate and ligand atoms, but these are used for book-keeping during training (see Section 3, not as features. We also compute ESM3 [4] embeddings of the residues and broadcast these to the residue's atom nodes; sequence and structure inputs were used when embedding proteins with ESM3.

Two geometric quantities are cached for each edge. First we compute a unit vector indicating the direction of the edge. The distance between the corresponding atoms is represented as the values of 16 radial basis functions. We use a Bessel basis on $[0, 8.0]$ Å with a smooth polynomial cutoff, so every feature decays to zero as an edge approaches the cutoff distance. We clamp distances below at $10^{-5}$ Å to avoid division by zero errors.

#### 2 Flow model architecture

##### 2.1 Message Passing on Atom Graphs

The generator is a message-passing graph network. Its input is the all-atom graph described in §1.6; its output is one "velocity" vector per water node, the displacement that carries that water along the flow (§3). Every node carries a feature vector, and the network refines these features over three rounds (layers) of message passing. In each layer, each node receives a message along every incoming edge, sums the incoming messages, and updates its own features from that sum. Successive layers let each node accumulate information from a widening graph neighborhood. After the final update layer, a readout layer maps each water node's features to its output velocity vector.

The graph for a protein has two node types: context (protein, ligand, symmetry mates) and water, and the message sent along an edge is computed by a function specific to that edge's type, with its own parameters. The reported models in the main text keep only protein–protein and protein-to-water edges, so water nodes are never the source of a message and are never connected to one another. A candidate water's features therefore depend only on its own position and its local protein environment. At a destination node the messages from its incoming edge types are summed and passed to an update function specific to that node type. Protein and water nodes carry separate update parameters.

Each node's features are a scalar–vector pair: $\nu = 256$ scalar channels that are invariant to rotation and $\mu = 64$ vector channels that rotate with the input structure. The scalar channels are initialized from an encoding of the node's identity (below). The node vector features are initialized to zero; they will be updated later during message passing, initially from the edge features as described in the next section.

##### 2.2 Message Function

The message and update functions use geometric vector perceptrons (GVPs, §2.4) [5] that update scalar–vector paired features on a node, keeping the scalar part invariant and the vector part equivariant (defined below). For an edge from source $i$ to destination $j$ the message is

$$m^{\rho}_{i\to j} = \mathrm{GVP}^{\rho}([s_i : e_{ij} : s_j], [\hat{u}_{i\to j} : V_i : V_j]), \tag{1}$$

where the superscript $\rho$ indexes the edge type (protein–protein or protein–water), each with its own GVP weights, $i \to j$ is a directed edge from source $i$ to destination $j$, and $[\cdot : \cdot]$ is concatenation. The GVP's scalar input is the concatenation of the source and destination scalars with the edge's radial features $e_{ij}$; its vector input is the concatenation of the edge unit vector $\hat{u}_{i\to j}$ with the source and destination vector channels $V_i$ and $V_j$. $V_i$ and $V_j$ are zero in the first layer, but since the edge unit vector always contributes one vector channel, they will be non-zero in subsequent layers. Each message function is a stack of two GVPs.

##### 2.3 Node Update Function

Within a block, each destination node aggregates its incoming messages by summing them, first within an edge type and then across the node's edge types, giving one aggregated message per node. The node's features are then refined in two residual sublayers, closely following [5]. First the aggregated message is added to the node's current features and the sum is layer-normalized. Then a node-update function transforms that result, and its output is in turn added back and layer-normalized. The update function is a stack of two GVPs that acts on the node's own scalar–vector pair alone, with no edge input; it is what mixes the newly received vector channels with the scalars inside a node and produces the features that the next block reads. Protein and water nodes carry separate message and update parameters, and dropout is applied at a channel level to the message and to the update before each residual add.

The message-passing backbone (message-passing and node update) has 256 scalar channels, 64 vector channels, and three message-passing blocks.

##### 2.4 Geometric Vector Perceptron

A vector-gated GVP (Algorithm 1 of [5]) acts on scalar channels $s \in \mathbb{R}^{\nu}$ and vector channels $V \in \mathbb{R}^{\mu\times 3}$:

$$\begin{aligned} V_h &= W_h V, & s_m &= W_s[\, s : \|V_h\|_2\,], \\ s' &= \sigma(s_m), & V' &= \sigma_g\big(W_g\, \sigma^{+}(s_m) + b_g\big) \odot W_v V_h. \end{aligned} \tag{2}$$

The $W_i$ are learned weight matrices, $V_h = W_h V$ is an intermediate set of $h$ vector channels, and $s_m$ is the pre-activation scalar. Here $\|\cdot\|_2$ is a row-wise $L_2$ norm of the vector features, $\odot$ is an element-wise product, and $[\cdot : \cdot]$ indicates concatenation. $\sigma$, $\sigma^{+}$, and $\sigma_g$ are non-linear "squashing" functions; following [5] we use a sigmoid and ReLU, respectively. Scalar channels depend on the vector features only through their norms, so they are rotation- and translation-invariant; vector channels are linearly recombined and gated by scalars, so they rotate with the input, i.e., they are equivariant. A GVP therefore returns invariant scalar features and equivariant vector features, and the per-water velocity read out at the end rotates with the input structure.

##### 2.5 Scalar Feature Encoders

The scalar channels are initialized by encoders that run once, before message passing layers. For context atoms (protein, ligand, symmetry mates), the one-hot element identity and the ESM3 residue embedding [4] are projected to a common width, normalized separately, concatenated, and passed through an MLP. Separate normalization keeps the much larger raw ESM3 scale from overwhelming the one-hot element channel. A symmetry mate protein atom inherits the embedding of its source ASU residue, and ligand atoms have their ESM3 embedding set to zero. The fused context features are then projected to the scalar backbone width by a linear layer with ReLU (implemented as a GVP with no vector input). Water nodes receive no ESM3 embedding. A separate encoder projects their 16-dimensional oxygen element one-hot to the scalar backbone width. In each case one GVP projects the encoded scalars to the backbone width. It is given no input vector channels, consistent with the zero-initialized node vectors above.

Before message passing, the flow-matching time point is broadcast to every node and concatenated to the node scalars, then passed through a small MLP, using separate weights for protein and water nodes. No later block receives the time directly. A final GVP returns one velocity vector per water node.

#### 3 Flow-matching training

Flow-matching attempts to learn a model to predict the linear optimal-transport probability path from prior point $x_0$ to the target $x_1$ (a deposited water), i.e., $x_t = (1-t)x_0 + tx_1$. The conditional velocity is $x_1 - x_0$ [10], giving the objective

$$\mathcal{L}(\theta) = \mathbb{E}_{t,(x_0,x_1)} \left\| v_\theta(x_t, t) - (x_1 - x_0) \right\|^2 , \tag{3}$$

with $t$ sampled uniformly per structure and squared error averaged over water nodes.

To sample initial water positions $x_0$, we draw an eligible anchor atom uniformly at random, then select a point inside a sphere centered on that atom. The direction is chosen uniformly on the unit sphere, and the radius is $RU^{1/3}$ for $U \sim \mathcal{U}(0,1)$, where $R = 8.0$, the same as the contact radius used in constructing the protein-water graph. In contact-aware graphs, eligible anchors exclude mate atoms but include ASU protein and retained ASU ligand atoms. Every prior point is therefore within the graph cutoff (8.0 Å) of an eligible ASU context atom.

The prior and deposited point sets are paired to form a coupling distribution $\pi(x_0, x_1)$ per structure by

$$\pi^\star = \arg\min_{\pi \in S_N} \sum_{i=1}^{N} \left\| x_0^{(i)} - x_1^{(\pi(i))} \right\|^2 , \tag{4}$$

solved exactly with the Hungarian-matching algorithm, a minimum-cost one-to-one bipartite matching method. This is the discrete optimal-transport assignment between the two complete point sets for one training example, following convention in [19, 16].

Training with both Small and Large datasets used AdamW [12] with learning rate $2 \times 10^{-3}$, weight decay $3 \times 10^{-2}$, unit-norm gradient clipping, and torch's automatic downcasting to `bfloat16`. The effective batch size is 16: four H100 GPUs with per-GPU batch one and four accumulation steps for Small, and eight H100 GPUs with per-GPU batch one and two accumulation steps for Large. Training uses 300 linear warmup steps and a 100-epoch cosine schedule within a 110-epoch budget.

A generative validation run is performed every two epochs on the validation structures using 50-step Euler integration, and sampling ratio of 1 water per residue. Checkpoints are ranked by a three-evaluation rolling mean of $0.85F_1 + 0.15\,\mathrm{AUC\text{-}PR}$ (metrics defined in section 7). The weights in the blend functions are hyperparameters. The selected symmetry-mate-aware checkpoints are epoch 74 for Small and epoch 98 for Large.

#### 4 Inference with the Flow Generator

Inference uses one preprocessed protein graph as input, and outputs a ranked set of water coordinates. First, inference with the flow generator draws $r = 8$ candidates per protein residue represented in the graph by default. Protein residues introduced from crystal contacts count toward this total; ligand atoms do not. Candidates are drawn from the same protein-anchored prior used during training (§3). WaterFlow then integrates the learned velocity field from $t_0$ to $t_1$ in 20 explicit-Euler steps [11]. At each step the model reads the current water coordinates and the fixed protein graph, predicts a displacement velocity for every water, and advances all waters by one time increment. The final coordinates form an intentionally redundant and dense candidate cloud. Several particles may converge on the same hydration site, and some may remain in unsupported positions.

#### 5 Confidence model

The confidence model has a similar overall architecture (see below) as the flow-matching generator, but is trained to predict whether a given water molecule is true or not. The confidence model does not update the coordinates. The input waters are generated by the flow-matching model described above. In case the flow-matching model predicts a water not within 8.0 Å of some protein atom, we connect it to the eight nearest protein atoms; this happens in less than 0.01% of cases. There is no one-to-one matching, so each target may have more than one predicted water mapped to it.

The confidence model uses the same GVP encoder architecture and hidden dimensions as the generator. Unlike the generator, we do not include a time input (since the confidence model is not a flow-matching model) and it predicts a scalar logit rather than a vector field. As in the generator, message passing moves information along protein-to-protein and protein-to-water atom edges. Since there are no water-to-water edges or water-to-protein edges, each candidate water's score is independent of other candidate waters, which is critical to the use of the model as scoring each water independently, not scoring the entire water configuration which would be much more complex.

Candidate water molecules at a distance $d$ from the nearest deposited water are assigned the smootherstep [15] target value

$$c(d) = \begin{cases} 1, & d \le r_{\text{in}}, \\ 1 - S\left(\dfrac{d - r_{\text{in}}}{r_{\text{out}} - r_{\text{in}}}\right), & r_{\text{in}} < d < r_{\text{out}}, \\ 0, & d \ge r_{\text{out}}, \end{cases} \quad S(u) = 6u^5 - 15u^4 + 10u^3, \quad r_{\text{in}} = 0.5\,\text{Å}, \quad r_{\text{out}} = 1.5\,\text{Å}. \tag{5}$$

Its midpoint is 1.0 Å, matching the primary acceptance radius we use in evaluation, but we note that it is not a calibrated probability.

SuperWater [7] constructs its labels using a similar geometric function. It finds the minimum Euclidean distance $d_{min}$ from each sampled candidate to any retained crystal-water oxygen. It maps that distance to

$$N(d_{min}) = \left(\frac{2}{1 + e^{-4d_{min}/\ln 2}} - 1\right)^2. \tag{6}$$

This function increases from 0 at $d_{min} = 0$ to 1 as the candidate moves farther from a crystal water. The SuperWater confidence network therefore regresses something like a normalised distance. At inference, it reports $1 - min(1, N_{pred})$ as a confidence.

The SuperWater target (Eqn. 6) has half-height near 0.31 Å and is already near 0.01 at 1.0 Å. To rectify this, WaterFlow's target uses two radii instead to define an exact high-score plateau through 0.5 Å, a smooth transition centered at 1.0 Å, and an exact zero beyond 1.5 Å. WaterFlow then applies binary cross-entropy with logits directly to this soft target. The sigmoid of the learned logit is the deployed high-is-good score, so no reversal is needed at inference.

Unlike SuperWater, we initialize the confidence model from WaterFlow's flow-matching candidate generator wherever possible. Training lasts 25 epochs with AdamW, learning rate $10^{-4}$, weight decay 0.01, 300 warmup steps, a cosine schedule, and effective batch size 16 on H100 GPUs. At most 1,500 randomly selected candidates from a structure are used in a training pass. While the confidence head is trained against a continuous target, we validate our model by treating it as a ranker. Each candidate is given a label, where it is counted as positive when its nearest ground truth water lies within 1.0 Å. We pool the confidence score and label pairs of every candidate in the validation set to compute metrics, rather than first computing metrics per protein and then averaging, and compute the area under the precision-recall curve (AUC-PR) to select the best checkpoint.

The retrained SuperWater confidence models use the score model's graph settings and frozen score checkpoint to generate their candidates, with 15 candidates per receptor residue and 20 reverse-diffusion steps. However, the saved configurations set `transfer_weights=false`: the confidence network shares the score model's tensor-product architecture, but training starts with freshly initialized weights. It is trained for 30 epochs with Adam, learning rate $10^{-3}$, zero weight decay, batch size 2, and a cosine schedule. We selected the best checkpoint as the one which minimizes validation MSE on the normalized-distance target.

#### 6 Evaluation metrics

Our primary metrics are precision and recall. However, since we predict *coordinates* of a certain number of water molecules, we need to define what is a "true positive". We do that by setting an *acceptance radius* $d$ which we set to either 0.5 or 1.0 Å. Precision is defined as the fraction of predicted waters with a target water center within the acceptance radius of its center. Recall is defined vice-versa, as the fraction of target waters with a predicted water center within the acceptance radius of its center.

Mathematically, for structure $i$, let $\{\hat{x}_{im}\}_{m=1}^{M_i}$ denote predicted positions and $\{x_{in}\}_{n=1}^{N_i}$ deposited positions. The precision and recall over all structures $i$ in some evaluation set are

$$P(d) = \frac{\sum_i \sum_m \mathbf{1}[\min_n \|\hat{x}_{im} - x_{in}\| \le d]}{\sum_i M_i}, \qquad R(d) = \frac{\sum_i \sum_n \mathbf{1}[\min_m \|\hat{x}_{im} - x_{in}\| \le d]}{\sum_i N_i}. \tag{7}$$

We frequently report the $F_1$ score which is the harmonic mean of precision and recall:

$$F_1(d) = \frac{2P(d)R(d)}{P(d)+R(d)}. \tag{8}$$

Note that our approach does not necessarily match the prediction and target waters one-to-one: several predictions may match the same deposited water, following the SuperWater evaluation convention [7].

Threshold sweeps produce post-clustering precision–recall curves. For WaterFlow, candidates below the threshold are removed before clustering (§7). The cutoffs are 0.05, 0.10, 0.20, ..., 0.90. SuperWater uses 0.02, 0.05, 0.10, 0.15, ..., 0.50. Its target falls to 0.5 by about 0.31 Å from a reference water and to about 0.01 by 1.0 Å, so its useful cutoffs are concentrated toward the lower end of $[0, 1]$. WaterFlow instead learns a sigmoid score from a target whose transition is centered at 1.0 Å. Equal numerical cutoffs would therefore represent different geometric error regimes, and the scores are not calibrated against one another. Both SuperWater and WaterFlow use one fixed candidate cloud and their own postprocessing code throughout the sweep.

##### 6.1 Evaluation at "parity"

Very often in this work we evaluate the model by setting thresholds (or densities) such that the number of predicted water molecules is equal to the number of target waters for a given input protein structure. We refer to this point as "parity". When we sweep score cutoffs or densities, the ratio of predicted to target water molecules may not actually be exactly 1, and so we linearly interpolate to report precision, recall, and $F_1$ scores. Furthermore, because the prediction-to-target mapping is not necessarily one-to-one, precision and recall need not be exactly equal at that point.

##### 6.2 Area Under the Precision-Recall Curve

We use area under the precision-recall curve (AUC–PR) for checkpoint selection during validation for the confidence model as well as the flow-matching generator, although they are computed differently in each case, and the flow-matching generator uses a linear combination of F1 at 1 Å and the AUC–PR. In the case of the generator, we sweep the matching tolerance for each predicted water, i.e. the threshold at which we compute precision and recall, over 50 values spaced uniformly on $[0.1, 3.0]$ Å. For the confidence model, each candidate water is assigned a binary label where the label is 1 if the water is within 1 Å of a ground truth water, and 0 otherwise. Candidates are then ranked by the predicted confidence score, and we compute the area under the precision-recall curve over a single global ranking pooled across all the waters of the whole validation split.

#### 7 Post-processing of generated water candidates: clustering and thresholding

Our flow-matching generative model is given more candidate water positions than can possibly be found around a protein. As described above, we use a confidence model to determine a score for each of these waters, which can be applied in one of two ways: simple thresholding (Section 7.2) or using it to rank candidates and taking the top $N$ (Section 7.3). Additionally a clustering routine (Section 7.1) collapses nearby waters into one centroid, so that very similar predictions become one. The complete algorithm given the scored candidates is given in Algorithm 1.

**Algorithm 1** Water-site selection

**Input:** candidates $\mathcal{C} = \{c_i\} = \{(s_i, p_i)\}$ with scores $s_i$ and positions $p_i$, radius $r$, thresholding rule `rule`, threshold $t$ or ratio $\rho$

**Output:** predicted water positions in descending score order

```
if rule is CONFIDENCE then
    discard any candidate c_i if s_i < t
end if
    sites ← GREEDYWATERCLUSTERS(C, r)                ▷ §7.1
if rule is DENSITY(ρ, n_res) then                     ▷ §7.3
    K ← round(ρ n_res)
    sites ← TOPK(sites, K)
end if
return sites
```

Note that only one of DENSITY and CONFIDENCE is chosen. Because the cutoff is applied after clustering rather than before it, a low-scoring candidate that would have been discarded under confidence thresholding can still join a cluster and shift its weighted coordinate, so a site retained by both rules need not sit at the same position. In general, there is no guarantee even that if the two methods yield the same number of final water predictions, that those predictions are the same sites.

##### 7.1 Clustering

Our clustering shown in Algorithm 2 uses a simple greedy approach. In the first pass, the highest scoring candidate water is selected and all other candidates within some distance $r$ are clustered with it. In our case we always set $r = 1.52$ Å, the van der Waal's radius of oxygen. Then the position of the cluster is computed (by subroutine ClusterMean, line 5) as a weighted mean of all candidate positions in the cluster, weighted by their scores. In the event that all scores in the cluster are zero, we use a simple arithmetic mean. Subroutine SuppressNeighbors (line 10 uses non-maximum suppression to remove additional overlapping clusters. Iterating through the `sites` in descending score order, it discards every lower-scoring site within $r$ of any cluster already kept.

**Algorithm 2** Greedy water site clustering GreedyWaterClusters

**Input:** candidates $\mathcal{C} = \{c_i\} = \{(s_i, p_i)\}$ with scores $s_i$ and positions $p_i$, radius $r$
**Output:** set `sites` water cluster centroids

1: `sites` $\leftarrow \varnothing$
2: **while** $\mathcal{C}$ is not empty **do** ▷ pass 1: absorb
3: $\quad i^* \leftarrow \text{argmax}_i\{s_i\}$, $c_i \in \mathcal{C}$
4: $\quad$ cluster $\mathcal{X} \leftarrow \{c_i \in \mathcal{C} \mid \|p_i - p_{i^*}\| < r\}$
5: $\quad$ position $\leftarrow$ ClusterMean($\mathcal{X}$)
6: $\quad$ score $\leftarrow \max_{(s_i,p_i)\in\mathcal{X}} s_i$
7: $\quad$ add (position, score) to `sites`
8: $\quad \mathcal{C} \leftarrow \mathcal{C} \setminus \mathcal{X}$
9: **end while**
10: sites $\leftarrow$ SuppressNeighbors(sites, $r$) ▷ pass 2

Clustering pass 1 produces water positions in descending score order, because the seed at line **??** is the highest-scoring *unassigned* candidate and each member of its cluster not previously assigned. SuppressNeighbors therefore does not need to sort and simply iterates through the list of water clusters `sites`; any score ties are therefore broken by input order.

Clustering pass 2 is not redundant with pass 1. Pass 1 tests distance to *candidate* water molecules. The weighted cluster means can leave two sites within $r$ of each other. Line 10 removes that residual duplication.

##### 7.2 Confidence thresholding

The standard path to predict waters accepts a minimum confidence $t$. Every candidate scoring below $t$ is discarded, and the remainder are consolidated as above. Thresholding occurs before any coordinates are averaged, so a rejected candidate can neither seed a cluster nor shift the position of an accepted one. The size of the output is set by the model's own confidence and therefore varies from structure to structure.

##### 7.3 Density thresholding

Density thresholding replaces the score threshold with a requested number of waters per protein residue in the asymmetric unit, defined by a ratio $\rho$. We determine the number of final water sites to retain as $K = \text{round}(\rho \times n_{\text{ASU}})$ where $n_{ASU}$ is the number of protein residues in the asymmetric unit. All candidates are clustered with no confidence cutoff using Algorithm 2. The resulting centroids are sorted by their scores assigned in line 6 of Algorithm 2, and the top $K$ are returned, or all available centroids if fewer exist. This path is useful when a downstream protocol needs a predictable output size.

##### 7.4 Structure output

The prediction script writes out a complete crystallographic model including both the protein and ligand atoms used as input, with the predicted water oxygen positions as described above. The retained water cluster centroids are returned to the input crystal frame (de-centered), merged into the input structure as solvent, and written in the input file's format alongside a plain-text table of scored coordinates.

**Frame.** The model works in a frame centered on the ASU protein-atom centroid (§1.6). That centroid is retained at graph construction and added back to every selected coordinate before anything is written, so the output is aligned to the input model with no further transformation. The predicted waters are the only new atoms. Protein and heteroatom coordinates pass through untouched.

**Solvent records.** Each retained centroid becomes one oxygen atom, named `O`, in its own `HOH` residue, flagged as a heteroatom and numbered sequentially. The waters are placed in a single chain whose identifier is chosen to be one not already present in the input, so as to not interfere with existing chains or residues. The occupancy for all predicted waters is set to 1.0. When the input carries $B$ factors, each water is assigned the mean $B$ factor of the input atoms within 5.0 Å of it, plus 10.0 Å$^2$. If no input atoms are within 5.0 Å, $B$ is set to the overall mean $B$ of the protein atoms plus the offset.

**Crystallographic metadata.** The unit cell is carried through from the input, and the space-group is read from the input and written explicitly. Output format follows the requested extension, mmCIF for `.cif` or `.mmcif` and PDB otherwise. A companion `_waters.txt` lists $x$, $y$, $z$ and the confidence score of every written water, so downstream code can re-rank or re-filter the output without re-reading the structure.

##### 7.5 Comparing the Density and Confidence rules

We compared the two filtering rules used in Algorithm 1 on the 698-structure holdout (§??). For each structure we recorded the number of retained waters per ASU protein residue under each rule and compared it with the deposited water density and the deposited resolution. The paired settings are $t \in \{0.30, 0.50, 0.75, 0.90\}$ against $\rho \in \{0.30, 0.60, 0.80, 1.00\}$. Where the analyses below require a single pair of settings rather than a sweep, they use $c = 0.50$ and $\rho = 0.60$, the pair chosen so that the two rules return comparable numbers of waters, and refer to it as the matched operating point.

Desnity thresholding does not vary with resolution. $\rho$ fixes waters per residue, hence this quantity scales with only the size of the protein. We therefore report the resolution relationship for the confidence rule and the deposited waters, and treat the independence of desnity thresholding from resolution or Wilson B-factor as a property of how it is defined rather stating the measurement.

#### 8 Data-scaling and ablation experiments

##### 8.1 Data scaling

To measure the impact of training-set size on precision and recall, we trained a set of models on nested sub-samples of the Large and Small training sets, from a few hundred structures up to the full sets. We train the generator and confidence model as a pair with the same training data. We then run inference with each trained pair of models on the 698-protein holdout set and compute $F_1$ at 1.0 Å with the predicted number of waters set to the target number of waters. The 1, 2 and 3% sub-samples are each retrained with different random seeds, from which we compute the error bars shown in Figure 3 of the main text; models trained with sub-samples of 10% and above are only trained once.

##### 8.2 Quality-filter ablation

To test how the training-time water filter (EDIAm $\geq 0.4$ and B-factor z-score $\leq 2.0$, §1) affects accuracy, we trained one model on the same dataset as our reported model (Large), but without the water quality filters, thereby keeping every deposited water. We compared it to the standard reported model with filtered waters in the training set on the 698-protein holdout set. Because the two return different numbers of waters at a given threshold, we compare them at matched water count, plotting $F_1$ at 1.0 Å acceptance radius against waters predicted per ASU residue, so that the confidence threshold does not confound our analysis. Both runs use the confidence thresholding rule, which is not the variable under test, and both are scored against the standard filtered target, which is the filtered experiment's own training target and therefore the conservative choice.

##### 8.3 Symmetry-mate ablation

To measure the contribution of symmetry-mate context (§1.5), we compare WaterFlow trained on Large with symmetry mates against an otherwise identical model trained and evaluated without symmetry mates, on the 698-protein holdout set. We report precision and recall over the confidence sweep and, to localize where the mates help, recall at parity binned by each deposited water's distance to the nearest symmetry-mate atom, in uniform 0.5 Å bins from 2.5 to 8.0 Å. Waters that bridge the asymmetric unit to its symmetry-related neighbors fall in the shortest-distance bins. The 2.0–2.5 Å bin is omitted. It holds 529 waters (compared to 14–28k in each other bin), and represents a near-clash regime in which both models are poor and the estimate is noisy.

### 9 Electron-density validation

Matching a prediction to a deposited water measures agreement with the atomic model. As discussed in the paper, it does not measure agreement with the underlying diffraction data. We therefore also score every predicted water against the deposited electron-density maps. Electron density maps still carry some bias from the atomic model, but represent a next step towards the raw data from the deposited atomic model.

#### 9.1 What is scored

Each measurement compares one set of predicted waters against one experimental map. The waters come from the 698-structure holdout (§1.2), predicted with the confidence threshdoling rule at threshold $t \in \{0.10, 0.30, 0.50, 0.75, 0.90\}$ and following the pipeline of Section 7 exactly, so candidates $c_i$ with scores $s_i < t$ are discarded before clustering and each threshold gives its own set of waters.

Predicted waters are returned to the crystal coordinate frame (§7.4), given occupancy 1.0, and assigned B-factors based on the rule in 7.4. That $B$ factor is not derived from the confidence score, and it does not affect EDIAm, which integrates the map over a sphere of fixed per-element radius and weights grid points by distance alone [14]. It does affect the $R_{\text{free}}$ test of Section 10, which is why that test also reports a regime in which water $B$ factors are refined.

#### 9.2 Map scoring

We measure density support with PDB-REDO [6] `density-fitness` 1.2.1, libpdb-redo 3.5.1 and Gemmi 0.7.3 [23]. We use only $2mF_o - DF_c$ (`FWT/PHWT`) and $mF_o - DF_c$ (`DELFWT/PHDELWT`) map coefficients from the deposited structure factor (MTZ) file. We do not calculate maps using any WaterFlow model, so every set of predicted waters for a given structure is scored against the same experimental map. We keep the program defaults, including X-ray scattering factors and a map-sampling factor of 1.5, and we pin these versions because recent libpdb-redo releases corrected the EDIA calculation and return different values from older builds.

The reported metric is EDIAm, the electron-density support at the water oxygen [14]; a larger EDIAm indicates closer agreement with the local map. A water is called well supported when its EDIAm is at least 0.8, the cutoff proposed by the paper that defines the metric.

#### 9.3 Matched and novel predictions

A prediction is *matched* when its nearest deposited water lies within 1.0 Å and *novel* otherwise; the same density or confidence rule and radius are used wherever predictions are split by category.

The distinction matters because of phase bias. Deposited waters contributed to the phases of the structure factors used to compute the deposited $2mF_o - DF_c$ map; as a result the map will favor deposited waters and potentially disfavor novel waters. We therefore report matched and novel EDIAm separately rather than pool them together. Furthermore we test novel predictions against the residual $mF_o - DF_c$ map as well.

#### 9.4 Density support across the confidence sweep

Both the EDIAm sweep and the residual-density experiments average statistics over all waters in the holdout dataset rather than averaging over the per-structure statistics. Thus each experiment shows the full per-water distribution at every threshold. At each confidence threshold $t$ we score every predicted water's EDIAm and compare that distribution to that of the deposited waters.

For novel predictions we also examine the $mF_o - DF_c$ "difference density" map. The deposited model has no water at novel sites, so a positive peak marks density that is unexplained by the deposited model. We also sample both maps at each novel prediction and at a displaced null/decoy. We include up to eight decoys per prediction, each displaced 2.5 Å from the novel water prediction, and retained only if it is not within 2.4 Å of any other non-water heavy atom. Decoys are drawn with rejection sampling for up to 48 attempts. We report densities in $\sigma$ (standard deviations of the map). We further report the fraction of novel predictions found in difference density $> +3\sigma$, against the same fraction for the decoys.

#### 10 Fixed-coordinate crystallographic cross-validation

This test asks whether adding the water improves the model's disagreement $R_{\text{free}}$ with diffraction observations (reflections) withheld from refinement for validation [3]. $R_{\text{free}}$ is essentially the unexplained variance in the observations. Lower $R_{\text{free}}$ is better.

We report this for one example structure. 4RKW, solved at 1.50 Å resolution. We also studied this protein in Sections 9 and 11. Three sets of predicted waters are compared, all built as described in §9 on the same deposited protein and ligand atoms: the deposited waters, predictions at confidence threshold $t = 0.50$, and predictions at density ratio $\rho = 0.60$ waters per ASU protein residue. The sets differ in size, so the deposited result is a reference point.

##### 10.1 Refinement regimes

Each water set is first rewritten with Phenix `pdbtools` to drop the deposited group-displacement (TLS) records, which `phenix.refine` rejects in these models. We then run `phenix.refine` [1] using one of three protocols, each starting from that same rewritten file in its own working directory.

`phenix.refine` does not optimize all parameters at once. It cycles through the ones the user asks it to refine, such as atomic positions, isotropic B-factors (displacement parameters), and then occupancies. It optimizes each in turn, with the other parameters held fixed, and re-fits the bulk-solvent and overall scale terms at the start of each minimization pass. One such pass is a macrocycle. Setting the number of macrocycles parameter to 0 results in no atomic parameter being refined.

The three protocols we use are are:

***rephase*** Zero refinement macrocycles. Coordinates, $B$ factors and occupancies are held fixed, and Phenix fits only the bulk-solvent and overall scale terms needed to compute $R$ factors. This scores the predicted positions exactly as placed, together with their placeholder $B$ factor.

***water-B*** Five macrocycles of per-atom $B$-factor refinement, with water oxygens restricted to isotropic $B$-factors and occupancies left out, so each water's $B$ factor is its only free parameter. All coordinates stay fixed. This removes the placeholder $B$-factor from the comparison and isolates the positions.

***full*** Five macrocycles with all coordinates and individual $B$-factors free, so the waters relax into the density instead of being scored where they were placed.

#### 11 Tests with predicted protein coordinates

WaterFlow normally receives an experimental protein structure. Here we test its sensitivity to replacing those coordinates with a predicted fold, on 4RKW, the single-chain entry we examined in Sections 9 and 10, against its 242 deposited waters, and 231 remaining waters targets after quality filters.

##### 11.1 Substituting the protein coordinates

For this comparison, we used ESMFold [9] to predict a structure from 4RKW's deposited sequence, using the `facebook/esmfold_v1` checkpoint through the HuggingFace transformers implementation of `EsmForProteinFolding`. 4RKW was chosen from the holdout set of proteins in particular because the deposited structure has no alternate conformations, so there is no ambiguity about which atomic coordinates to choose. The two inference runs of the crystal structure and predicted structure have the same node features, ESM embeddings, water targets, number of water candidates generated, ASU residue count, and random seed.

##### 11.2 Selection and evaluation

Each candidate cloud from the predicted and the crystal structure are processed the same way as described in 7. The confidence thresholds are swept from 0.1 to 0.9, and the density thresholds are swept from 0.3 to 1.2 waters per ASU residue, both in steps of 0.1.

Specific case studies to show lost/gained waters in specific sites for 4RKW use either a confidence threshold $t = 0.5$ or density ratio $\rho = 0.6$.

#### 12 Comparison with SuperWater

SuperWater combines a score-based diffusion model, an $SE(3)$-equivariant tensor-product network, and ESM-2 embeddings, followed by a confidence model [7, 9], following a similar candidate generation followed by confidence scoring mechanism that we implement for WaterFlow. The published evaluation used random data splits. For this comparison, we retrained the SuperWater score models on the nested Small and Large training sets. The associated confidence models use these score checkpoints to generate candidates, but their weights are initialized and trained separately, as was done originally for SuperWater.

##### 12.1 Retraining configuration

The retrained SuperWater generator models use an all-atom representation with 24 scalar and 6 vector channels and three convolutional layers. Training uses a learning rate of $10^{-3}$, no weight decay, dropout 0.1, a cosine schedule, and an exponential moving average of the weights with decay 0.999. Training is done with `torch.DistributedDataParallel` across 8 H100 80GB GPUs at a per-GPU batch of 8, for an effective batch of 64, and runs to a maximum of 150 epochs with early stopping on validation performance.

For both training the generators and generating candidate waters, SuperWater uses a ratio of 15 waters per residue, unchanged from the original implementation. The confidence model for SuperWater uses the nearest-water normalized-distance regression in Eq. 6. Inference with SuperWater's prediction pipeline also uses a sampling ratio of 15 candidate waters per residue, and 20 reverse diffusion steps for the diffusion-based generator.

We train the models on both the SMALL and LARGE datasets, and report the model with the higher performance on F1 at parity at 1.0 Å.

##### 12.2 Shared evaluation conditions

WaterFlow and our re-trained checkpoints of SuperWater use the same training splits. We add EDIAm and B-factor filters to the SuperWater dataset processing. We also similarly use 0.5 and 1.0 Å acceptance radii, nearest-neighbor metric, and aggregate parity definition (§6).

The pre-processed datasets that are used to train each model contain slightly different number of ground-truth waters, based on how each codebase processes the input training data, so we score each model against its own processed set of ground-truth water.

It is also important to note that the clustering rules differ between WaterFlow and SuperWater. WaterFlow uses a 1.52 Å radius in both passes of clustering. The reported SuperWater analysis uses 1.52 Å for absorption and 2.2 Å for its second clash-removal pass.

#### 13 Clustering analysis of water molecules

Water molecules more than 4.0 Å away from the protein are removed from the analysis. The symmetry-equivalent water position yielding the closest distance to the protein is identified before applying this cutoff. Structures are aligned before clustering, and the alignment reference structure is selected to be the highest-resolution isomorphous structure, which is 7der for the 65 hen egg-white lysozyme (HEWL) structures, 5r32 for the 907 endothiapepsin (EAP) structures, and 3ks3 for the 937 carbonic anhydrase (CA) structures.

Water clustering is performed using HDBSCAN [13], which is also used by other packages that analyze conserved water molecules [2]. Code for pre-processing, clustering, analysis, and visualization is available here: https://github.com/diff-use/consensus_water_analysis/tree/v0.1.0. We performed hyperparameter grid-search over the Cartesian product of `min_cluster_size` and `min_samples` values. `min_cluster_size` controls the detection floor in clustering; we tried {5, 10, 15, 20}. `min_samples` controls the local density of the samples; we tried values of {2, 3, 5, 10, 15, 20} but capped it at `min_cluster_size`. Each hyperparameter combination of `min_cluster_size` and `min_samples` is ranked by two metrics. The first metric is DBCV [66], which scores the separation of clusters. The second metric scores the stability of clusters under each `min_cluster_size` by averaging the best Jaccard overlap of clusters obtained using different `min_samples`. More stable clusters have greater overlap for different values of `min_samples`. For each hyperparameter combination, the worse ranking of the two metrics is kept as its final ranking. We find the best clustering by this final ranking. The best clustering for each dataset is obtained with `min_cluster_size` and `min_samples` values shown in Supplementary Table 5.

#### References


[1] Pavel V. Afonine, Ralf W. Grosse-Kunstleve, Nathaniel Echols, Jeffrey J. Headd, Nigel W. Moriarty, Marat Mustyakimov, Thomas C. Terwilliger, Alexandre Urzhumtsev, Peter H. Zwart, and Paul D.

Adams. Towards automated crystallographic structure refinement with phenix.refine. *Acta Crystallographica Section D*, 68(4):352–367, 2012.

[2] Alfie-Louise R. Brownless, Travis Harrison-Rawn, and Shina C. L. Kamerlin. Watcon: A python tool for analysis of conserved water networks across protein families. *JACS Au*, 5(12):6379–6395, 11 2025.

[3] Axel T. Brünger. Free r value: a novel statistical quantity for assessing the accuracy of crystal structures. *Nature*, 355(6359):472–475, 1992.

[4] T. Hayes et al. Simulating 500 million years of evolution with a language model. *Science*, 387(6736):850–858, 2025.

[5] B. Jing, S. Eismann, P. Suriana, R. J. L. Townshend, and R. Dror. Learning from protein structure with geometric vector perceptrons. In *ICLR*, 2021.

[6] R. P. Joosten, F. Long, G. N. Murshudov, and A. Perrakis. The pdb_redo server for macromolecular structure model optimization. *IUCrJ*, 1(4):213–220, 2014.

[7] Xiaohan Kuang, Yunchao Lance Liu, Xiaobo Lin, Jesse Spencer-Smith, Tyler Derr, Yinghao Wu, Hans Bitter, Yongbo Hu, Jens Meiler, and Zhaoqian Su. Superwater as a generative AI framework to predict water molecule positions on protein structures. *Communications Chemistry*, 8(1):397, 2025.

[8] P. Kunzmann and K. Hamacher. Biotite: a unifying open source computational biology framework in python. *BMC Bioinformatics*, 19:346, 2018.

[9] Z. Lin et al. Evolutionary-scale prediction of atomic-level protein structure with a language model. *Science*, 379(6637):1123–1130, 2023.

[10] Y. Lipman, R. T. Q. Chen, H. Ben-Hamu, M. Nickel, and M. Le. Flow matching for generative modeling. In *ICLR*, 2023.

[11] Y. Lipman et al. Flow matching guide and code. *arXiv preprint arXiv:2412.06264*, 2024.

[12] I. Loshchilov and F. Hutter. Decoupled weight decay regularization. In *ICLR*, 2019.

[13] Leland McInnes, John Healy, and Steve Astels. hdbscan: Hierarchical density based clustering. *Journal of Open Source Software*, 2(11):205, 2017.

[14] A. Meyder, E. Nittinger, G. Lange, R. Klein, and M. Rarey. Estimating electron density support for individual atoms and molecular fragments in x-ray structures. *J. Chem. Inf. Model.*, 57(10):2437–2447, 2017.

[15] K. Perlin. Improving noise. *ACM Transactions on Graphics*, 21(3):681–682, 2002.

[16] A.-A. Pooladian et al. Multisample flow matching: straightening flows with minibatch couplings. In *ICML*, 2023.

[17] Schr
"odinger, LLC. The pymol molecular graphics system, 2024.

[18] M. Steinegger and J. S
"oding. Mmseqs2 enables sensitive protein sequence searching for the analysis of massive data sets. *Nat. Biotechnol.*, 35:1026–1028, 2017.
